# Seasonal and spatial controls on N_2_O concentrations and emissions in low-nitrogen estuaries: Evidence from three tropical systems

**DOI:** 10.1101/650705

**Authors:** Rachel Murray, Dirk Erler, Judith Rosentreter, Naomi Wells, Bradley Eyre

## Abstract

Estuarine N_2_O emissions contribute to the atmospheric N_2_O budget, but little is known about estuary N_2_O fluxes under low dissolved inorganic nitrogen (DIN) conditions. We present high-resolution spatial surveys of N_2_O concentrations and water-air fluxes in three low-DIN (NO_3_ ^−^ < 30 *µ*mol L^−1^) tropical estuaries in Queensland, Australia (Johnstone River, Fitzroy River, Constant Creek) during consecutive wet and dry seasons. Constant Creek had the lowest concentrations of dissolved inorganic nitrogen (DIN; 0.01 to 5.4 *µ*mol L^−1^ of NO_3_ ^−^ and 0.09 to 13.6 *µ*mol L^−1^ of NH_4_^+^) and N_2_O (93–132% saturation), and associated lowest N_2_O emissions (– 1.4 to 8.4 *µ*mol m^−2^ d^−1^) in both seasons. The other two estuaries exhibited higher DIN inputs and higher N_2_O emissions. The Johnstone River Estuary had the highest N_2_O concentrations (97–245% saturation) and emissions (– 0.03 to 25.7 *µ*mol m^−2^ d^−1^), driven by groundwater inputs from upstream sources, with increased N_2_O input in the wet season. In the Fitzroy River Estuary, N_2_O concentrations (100–204% saturation) and emissions (0.03–19.5 *µ*mol m^−2^ d^−1^) were associated with wastewater inputs, which had a larger effect during the dry season and were diluted during the wet season. Overall N_2_O emissions from the three tropical estuaries were low compared to previous studies, and at times water-air N_2_O fluxes were actually negative, indicating that N_2_O consumption occurred. Low water column NO_3_ ^−^ concentration (i.e. < 5 *µ*mol L^−1^) appears to promote negative water-air N_2_O fluxes in estuary environments; considering the number of estuaries and mangrove creeks where DIN falls below this threshold, negative water-air N_2_O fluxes are likely common.

## 1. Introduction

Humans have greatly altered the nitrogen cycle in estuarine and coastal waters through excess nitrogen loads from agriculture, wastewater and urban runoff (Howarth, 2008; Vitousek et al., 1997). Nitrogen over-enrichment drives the production of excess organic matter (eutrophication), which is one of the greatest threats to coastal ecosystems worldwide (Howarth and Marino, 2006; de Jonge et al., 2002). Nitrogen is biologically processed in estuaries via many pathways including assimilatory uptake and microbially-mediated redox-regulated processes (e.g. Salk et al., 2017; An and Gardner, 2002; Cook et al., 2004; Eyre et al., 2016). Some of these pathways produce nitrous oxide (N_2_O), which is a potent greenhouse gas that contributes about 8% of the radiative ‘greenhouse gas’ effect (Bouwman et al., 1995; Stocker et al., 2013) and causes ozone depletion in the stratosphere (Ravishankara et al., 2009).

In coastal areas, the most common biological pathways of N_2_O production are nitrification and denitrification. Denitrification is the reduction of NO_3_ ^−^ to NO_2_ ^−^, N_2_O, and N_2_ (Firestone et al., 1980; Firestone and Davidson, 1989). Typically, heterotrophic microbes are responsible for denitrification, however autotrophic ammonia-oxidising bacteria can reduce NO_2_ ^−^ to N_2_O and N_2_ via ‘nitrifier-denitrification.’ In contrast with denitrification, nitrification involves oxidation of NH_4_^+^ to N_2_O, NO_2_ ^−^, and NO_3_ ^−^ (Arp et al., 2002; Belser and Schmidt, 1978). Denitrification occurs in anoxic environments, such as anoxic layers of soil or sediment (Davidson and Swank, 1986; Firestone et al., 1979; Thomas et al., 1994), while nitrification occurs in both sediment (Jenkins and Kemp, 1984; Usui et al., 2001) and the water column (Damashek et al., 2016; Iriarte et al., 1998; Somville, 1984), where O_2_ is available.

High concentrations of nitrate (NO_3_ ^−^) in estuarine waters are linked to increased N_2_O water-air emissions (Murray et al., 2015), although this may be attenuated by the estuarine flushing time (Wells et al., 2018). High NH_4_^+^ concentrations are also known to enhance N_2_O production (De Wilde and De Bie, 2000). Most studies of N_2_O in tidal aquatic environments have been undertaken in disturbed estuaries with high DIN concentrations (up to 500 *µ*mol L^−1^) (Murray et al., 2015). Additionally, most N_2_O water concentration sampling has been undertaken in temperate or higher-latitude estuaries, with only 11 water-air N_2_O emission studies in estuaries located between 25°S and 25°N (Table 1). Waters low in dissolved inorganic nitrogen (DIN) can be a net sink for N_2_O, particularly in small mangrove-lined creeks (Erler et al., 2015; Maher et al., 2016; Wells et al., 2018; Murray et al., 2018). N_2_O undersaturation may be prevalent in mangrove creeks, but the extent and seasonal dynamics of N_2_O undersaturation are not well understood due to the limited number of studies and lack of seasonal sampling.

**Table 1:**
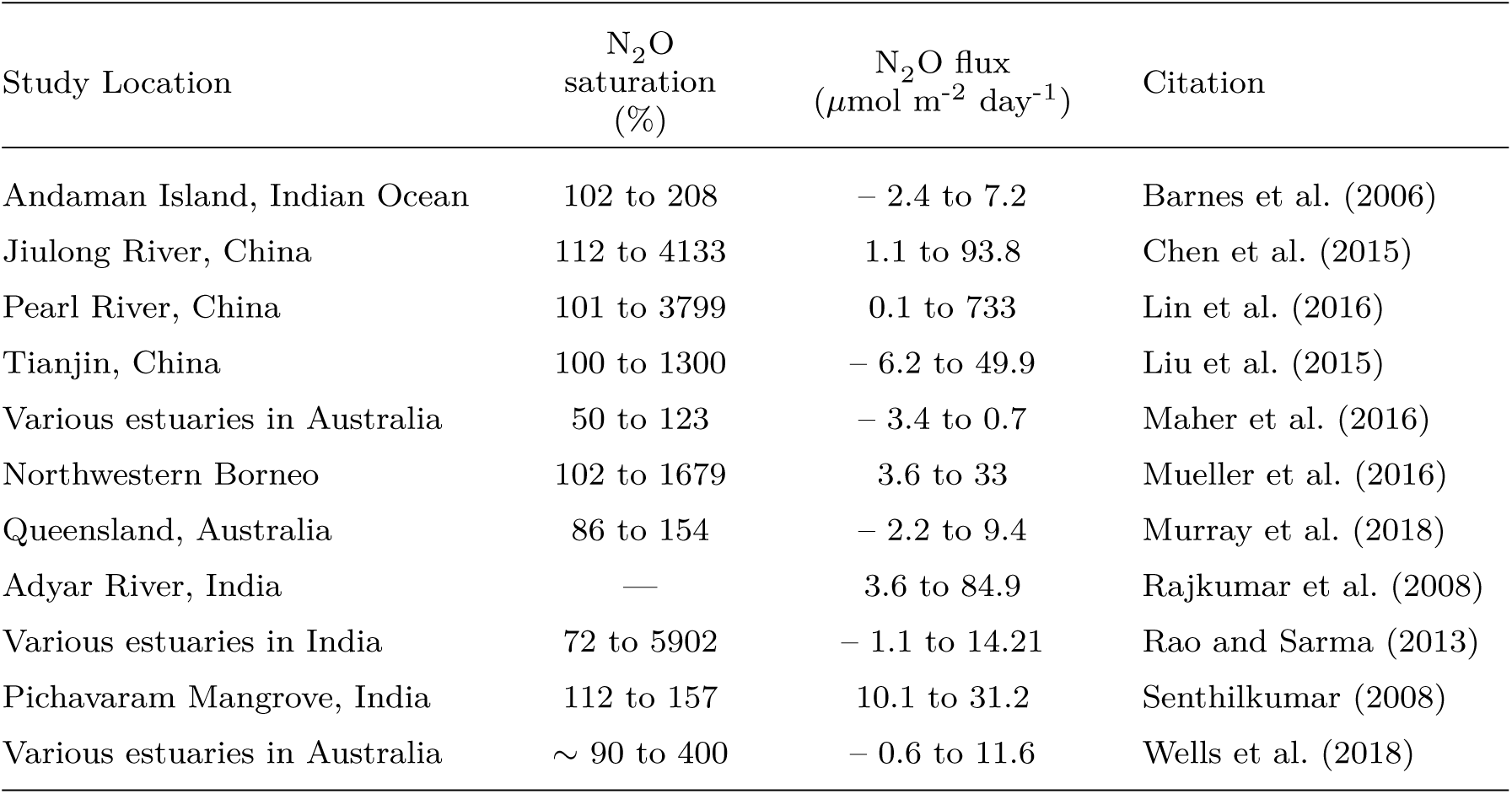
Range of N_2_O concentrations and water-air N_2_O fluxes in tropical estuaries.

| Study Location | $\text{N}_2\text{O}$<br>saturation<br>(%) | $\text{N}_2\text{O}$ flux<br>($\mu\text{mol m}^{-2} \text{ day}^{-1}$) | Citation |
| --- | --- | --- | --- |
| Andaman Island, Indian Ocean | 102 to 208 | – 2.4 to 7.2 | Barnes et al. (2006) |
| Jiulong River, China | 112 to 4133 | 1.1 to 93.8 | Chen et al. (2015) |
| Pearl River, China | 101 to 3799 | 0.1 to 733 | Lin et al. (2016) |
| Tianjin, China | 100 to 1300 | – 6.2 to 49.9 | Liu et al. (2015) |
| Various estuaries in Australia | 50 to 123 | – 3.4 to 0.7 | Maher et al. (2016) |
| Northwestern Borneo | 102 to 1679 | 3.6 to 33 | Mueller et al. (2016) |
| Queensland, Australia | 86 to 154 | – 2.2 to 9.4 | Murray et al. (2018) |
| Adyar River, India | — | 3.6 to 84.9 | Rajkumar et al. (2008) |
| Various estuaries in India | 72 to 5902 | – 1.1 to 14.21 | Rao and Sarma (2013) |
| Pichavaram Mangrove, India | 112 to 157 | 10.1 to 31.2 | Senthilkumar (2008) |
| Various estuaries in Australia | ~ 90 to 400 | – 0.6 to 11.6 | Wells et al. (2018) |

There can be significant variability in water-air N_2_O fluxes across larger estuaries due to N_2_O exchange, groundwater input, and benthic/water-column microbial activity. Large spatial N_2_O concentration gradients have been observed at ‘hot spots’ of N_2_O influx (Mueller et al., 2016; Wong et al., 2013). However, most estuarine N_2_O surveys employ discrete sampling at coarse resolution, which may not capture the full range of variation. Continuous sampling, employing real-time N_2_O concentration monitoring, is emerging as a way of investigating spatial variability in estuaries at a very fine scale (Bange et al., 1998; Brase et al., 2016; Mueller et al., 2016; Wells et al., 2018), however there has only been one such sampling campaign in a tropical estuary (Mueller et al., 2016).

In the current study, we present real-time N_2_O concentration measurements and interpret the drivers of N_2_O production and loss in three Queensland estuaries during the wet and dry seasons, focusing on catchments which are only slightly to moderately affected by anthropogenic disturbance. We hypothesise that in such environments, water-air N_2_O fluxes should be positive and increase with DIN concentration, however N_2_O may be undersaturated at very low DIN concentrations. The DIN–N_2_O relationship may provide some indication of the biogeochemical conditions under which a tropical estuary can transition from a net sink to a net source of N_2_O.

## 2. Study Locations

Surveys were conducted during March (wet season) and September (dry season) in northern Queensland, Australia, along the lengths of three estuaries: the Fitzroy River Estuary, the Johnstone River Estuary, and Constant Creek Estuary (Figure 1). These are wet and dry tropical estuaries where most of the rainfall occurs in summer, with a mean annual rainfall of 3152 mm in the Johnstone catchment, 1705 mm in the Constant Creek catchment, and 646 mm in the Fitzroy catchment (Queensland Department of Environment and Science). In the Johnstone and Fitzroy estuaries there is a strong seasonality in freshwater supply, as can be seen in data from upstream river gauges (Figure 2). Under high flows, salt water can flush fully to the mouth and during the dry season these systems can become hypersaline due to high evaporation and low freshwater input (Eyre, 1995, 1994).

**Figure 1:**
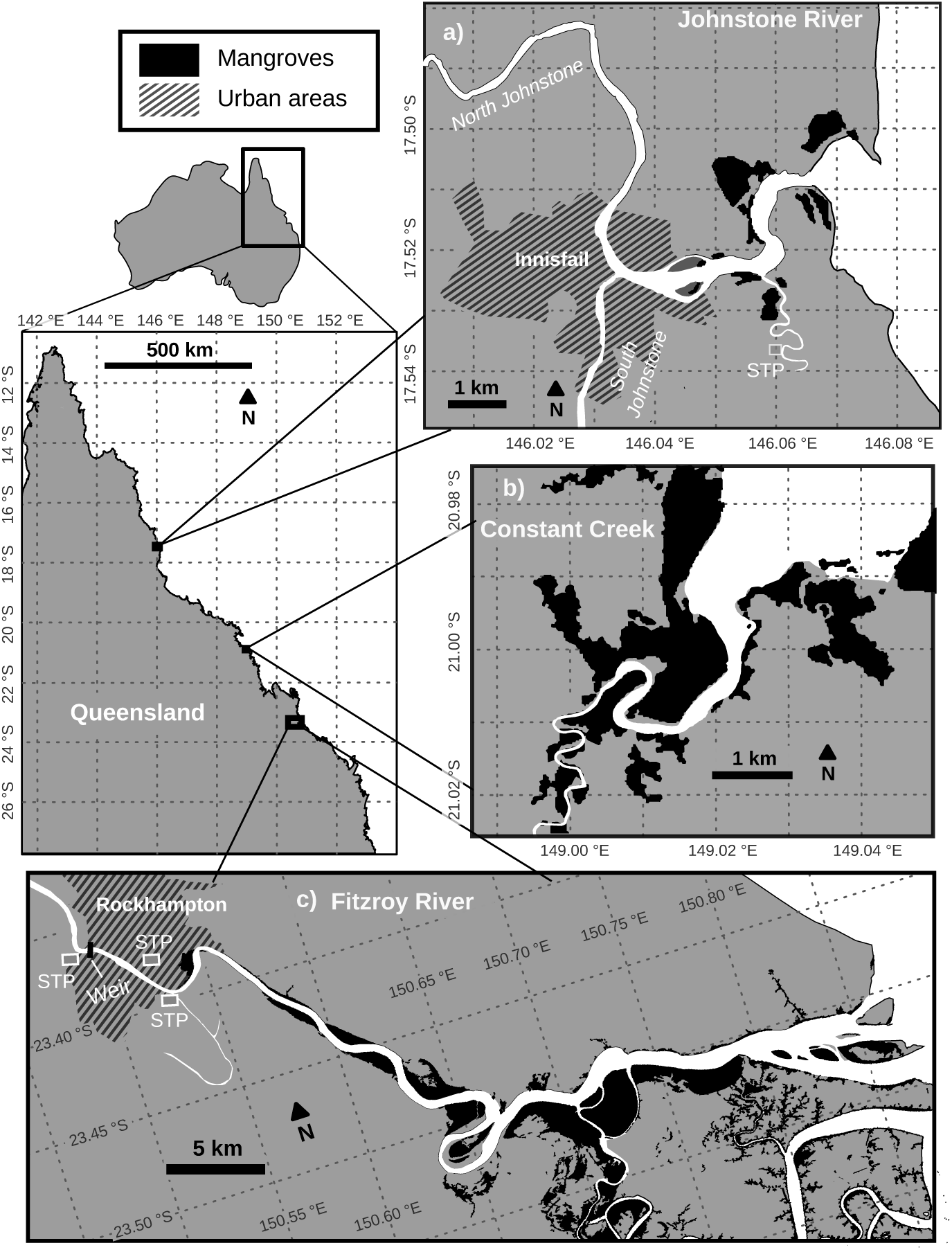
The three estuaries surveyed along the coast of Queensland, Australia: a) the Johnstone River Estuary, b) Constant Creek, and c) the Fitzroy River Estuary.

**Figure 2:**
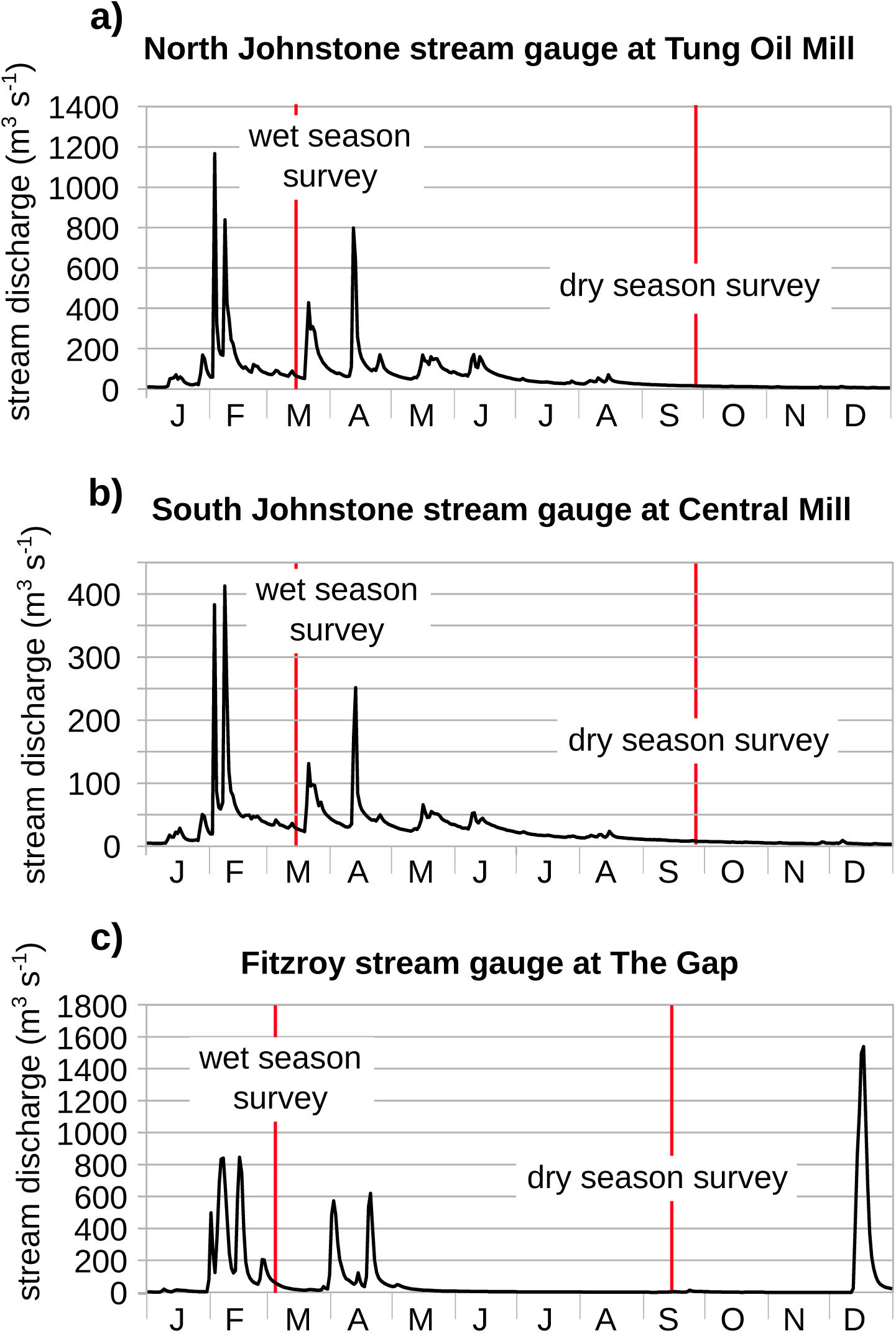
Stream discharge (m^3^ s^−1^) measurements from three tributaries upstream of the Johnstone and Fitzroy Rivers, at the locations of (a) ‘Tung Oil’ upstream of the North Johnstone River, (b) ‘Central Mill’ upstream of the South Johnstone River, and (c) ‘The Gap’ upstream of the Fitzroy River.

The Johnstone River is furthest north, at 17.5°S latitude, draining a catchment of 2320 m^2^. The mouth of the Johnstone River is just 5 km east of the city of Innisfail, QLD, which sits at the confluence of two major tributaries: the North Johnstone River and the South Johnstone River. In this catchment there is a sewage treatment plant located along a mangrove-lined creek, which empties into the Johnstone River about 2.5 km upstream of the mouth of the estuary (Figure 1a). The major land uses are banana and sugarcane cultivation. Constant Creek is located north of Mackay, QLD at about 21°S latitude, lined by mangroves and draining sugarcane cultivation and grazing lands (Figure 1b). The mouth of the Fitzroy River is located at about 23°S latitude, draining a catchment of 142,570 km^2^, and emptying into the ocean downstream of Rock-hampton, QLD. In the Fitzroy catchment, grazing and production forestry are the most common land uses. Several sewage treatment plants release effluent into the estuary just downstream of Rockhampton, near the upstream end of the estuary (Figure 1c).

## 3. Methods

### 3.1 Sample Collection and Analysis

Spatial surveys were performed on board a small vessel, starting at high tide at the mouth of each estuary and then moving upstream to the head of the estuary (Eyre, 2000). In the case of the Johnstone River estuary, the South Johnstone tributary was surveyed for about 1 km before a return trip to the fork in the river and a survey of the North Johnstone tributary. Water was continuously sampled from just below the water line through a water pump and hose, which were suspended from the side of the boat. This water was then fed into a shower-head style equilibrator to allow any dissolved gases to come into equilibrium between the water and a head-space of air (Johnson, 1999).

The head-space was vented into a second equilibrator head-space, which was itself vented into the ambient air. This assured there would be no pressure build-up in the equilibrator, while minimising atmospheric contamination of the sample. The head-space in the primary equilibrator was connected by 4 mm Bev-A-line tubing into a Picarro cavity ring-down spectrometer (CRDS) instrument (details described below). A Drierite^™^ column was connected to this tubing, between the equilibrator and the analyser inlet, to reduce humidity in the incoming air stream and minimise infrared interference from H_2_O. The analyser outlet was connected back to the equilibrator so that the CRDS and equilibrator formed a loop through which air continuously flowed. Initial experimentation revealed that there was a small lag (∼ 5 min) between water sampling and full equilibration of the water with the head-space.

Two different CRDS instruments were used to measure N_2_O concentration. In the wet season (March), the Picarro G5101-I isotopic N_2_O analyser (Picarro Inc., Santa Clara, CA. USA) was used (Erler et al., 2015). In the dry season (September) technical problems prevented the use of this device, so a G2308 Picarro was used. Both of these detect N_2_O by measuring the ‘ring-down’ time of infrared light, however there is a difference in precision; The G5101-I has a 1-minute 1-*σ* precision of ∼ 0.3 ppbv while the G2308 is less precise (∼ 8 ppbv). In both seasons a 350 ppbv N_2_O standard gas mixture was analysed to check for instrument drift before and after each deployment, and no significant instrument drift was recorded.

In addition to continuous N_2_O measurements, discrete water samples were collected about every 2 ppt of salinity along the length of the survey. These samples were filtered (0.45 *µ*m, Whatman GF/F), stored in 10 mL polycarbonate vials, and frozen for later DIN and *δ* ^15^N-DIN analysis. Salinity, temperature and luminescent dissolved oxygen (LDO) were measured every 5 minutes using a Hydrolab DS5X Sonde (AquaLab), which was suspended from the side of the vessel, just below the water line. This sonde was re-calibrated for salinity before and after each survey. ^222^Rn, a groundwater tracer, was measured every 10 minutes by a radon-in-air analyser (RAD7, Durridge) connected by a loop of tubing to the shower-head exchanger (Santos and Eyre, 2011). ^222^Rn data was only collected in the dry season.

Wind speed, required for N_2_O water-air flux calculations, was measured using an anemometer and augmented with 1-minute wind speed data procured from three Bureau of Meteorology (BOM) stations — Rockhampton (station number 039083), South Johnstone (032037), and MacKay (033045) (near Constant Creek). Depth datapoints came from a Garmin GPS unit on-board the boat. Discharge data (m^2^ s^−1^ of stream flow) came from two stream gauges upstream of the Johnstone River (stream gauge numbers 112004A and 112101B), and one stream gauge upstream of the Fitzroy River (stream gauge number 130005A) which were obtained from the Water Monitoring Information Portal of the Queensland Department of Natural Resources and Mines.

Discrete water samples were analysed for nitrate (NO_3_ ^−^), dissolved organic nitrogen (DON), and ammonium (NH_4_^+^) on a Lachat-flow injection device (see Eyre and Pont (2003); Eyre et al. (2011) for methods, errors and detection limits). Before *δ* ^15^N analysis, NO_3_ ^−^ was converted to N_2_O by denitrifying bacteria following the method of Sigman et al. (2001). The resulting N_2_O was then stripped from the solute by helium sparging and then analysed in a Thermo Fisher GasBench II which fed into to a Thermo Delta V Plus IRMS.

### 3.2. Calculations

The dissolved N_2_O and saturation values were calculated based on the solubility equations of Weiss and Price (Weiss and Price, 1980), and apparent oxygen utilisation (AOU) was derived from LDO using the solubility equations of (Benson and Krause, 1984). Excess N_2_O (ΔN_2_O) was calculated as the difference between measured concentration of N_2_O (in nmol L^−1^) and the theoretical concentration at 100% saturation, which we refer to as the “background” concentration:

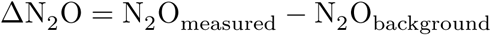

There are many parameterisations available for estimating water-to-air fluxes. The commonly-cited options can produce very different results (Musenze et al., 2014; Rosentreter et al., 2017), so fluxes were calculated using 6 different parameterisations (equations in Appendix Table A1). The mean fluxes presented in this study represent the average of all 6 parameterisations, and the ranges represent the full range of all calculated values. Gas fluxes were calculated at 1-minute intervals, and any missing temperature, salinity, or wind data was linearly interpolated from the nearest measured values. The resulting water-air N_2_O fluxes then were extrapolated over the area of the main channel of each estuary. To calculate the average area-weighted estuary water-to-air flux, it was first necessary to define a shape-file outlining the areal extent of each transect. This was done by manually digitising Landsat 8 satellite images for each estuary using QGIS software. After this step, the 1-minute-average water-air flux values (in *µ*mol N_2_O m^−2^ d^−1^) were extrapolated over the water surface area at ∼ 3 m resolution, using an inverse cost function – specifically, the GRASS GIS ‘v.surf.icw’ function (developed by Hamish Bowman) was used for this purpose. Finally, the area-weighted values were calculated as the average N_2_O flux of all of the pixels in the resulting geoTIFF raster image.

Once the total water-air N_2_O flux (*µ*mol d^−1^) was calculated, it was compared with various inputs into the estuary, such as the total DIN load (the DIN concentration of freshwater multiplied by the total freshwater discharge into each estuary) and the load of NO_3_ ^−^ and NH_4_^+^. Additionally, the depth (m), salinity, and N_2_O concentration (nmol L^−1^) datasets were extrapolated using the same method as the water-air fluxes – the inverse-cost function – over the same area in order to calculate the total water volume of each estuary and the total volume of freshwater. The freshwater fraction was calculated using the QGIS ‘raster calculator’ function at each pixel of the ‘depth’ geoTIFF as:

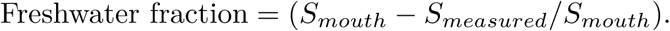

Where S_mouth_ is the salinity at the mouth of the estuary (the oceanic salinity) and S_measured_ is the salinity as measured (or extrapolated from measured values) at each point. The freshwater fraction values were multiplied by the depth and surface area at each pixel as:

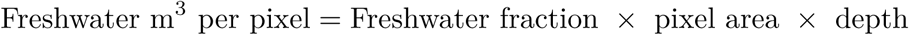

The sum of all values in the resulting raster image, calculated using the ‘raster statistics’ function in QGIS, was then the total amount (in m^3^) of freshwater in the estuary. This information, along with the freshwater discharge data, was required to calculate the freshwater flushing time using the fraction of freshwater method (Kennish, 1986), where the total freshwater volume is divided by the freshwater discharge at the upstream gauge. Instead of using the discharge on the day previous to sampling, however, the discharge of preceding days was summed until the total freshwater volume was reached (Eyre, 2000; Kaul and Froelich, 1984). The exception was at Constant Creek, where freshwater residence time was not calculated due to the absence of upstream discharge data.

The discharge data and total N_2_O flux data were used to estimate the contribution of ventilation of river N_2_O to the total N_2_O flux within each estuary (modified from Abril et al., 2000; Borges and Abril, 2011). This value represents the proportion of the positive water-air N_2_O flux which would be needed to offset all of the excess dissolved N_2_O (ΔN_2_O) delivered by the river at the upstream end of the estuary channel. It is possible for this proportion to be greater than 100% in estuaries where the water-air N_2_O flux is too low to fully account for all incoming dissolved N_2_O – i.e. in estuaries where freshwater-derived N_2_O is exported to the ocean.

Independently, the theoretical ventilation time (the time required to equilibrate the entire water column with overlying air) was estimated as the ratio between the average depth and average gas transfer velocity (Bender et al., 2011; De Wilde and De Bie, 2000; Reuer et al., 2007). In addition, where N_2_O concentrations fell below the conservative salinity mixing line, the amount of N_2_O removed from the water column 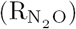, was calculated as the sum of the deviations between observed N_2_O concentrations and expected N_2_O concentrations at each point.

For consistency, the area of each transect was defined as the maximum area that had sampling coverage for both seasons. In the Fitzroy, the survey extended from the mouth of the estuary to a point just downstream of the Rockhampton weir. In the Johnstone, the sampling vessel traveled up the South and North Johnstone for a few km. In Constant Creek the surveyed area extended upstream to a bridge just inland of the boat ramp.

## 4. Results

### 4.1. Johnstone River Estuary

Nitrous oxide concentrations in the Johnstone River Estuary ranged from 132 to 245% saturation in the wet season and from 97 to 228% saturation in the dry season (Table 2; Figure 3e,f). Water-air N_2_O fluxes were higher than in the Fitzroy River Estuary and Constant Creek, with a wide range of 0 to 47.5 *µ*mol m^−2^ d^−1^ over the salinity gradient in the wet season (Table 2; Figure 3g,h) and an average area-weighted water-air flux of 5.6 *µ*mol m^−2^ d^−1^ (Table 2). The range over the salinity gradient in the dry season was – 0.3 to 28.8 *µ*mol m^−2^ d^−1^ and the average area-weighted water-air flux was 3.6 *µ*mol m^−2^ d^−1^. Nitrous oxide concentrations were generally highest at the freshwater end-member, particularly in the South Johnstone (Figure 4a,b), where N_2_O saturations reached as high as 245% in the wet season and 225% in the dry season. In the North Johnstone, N_2_O reached a maximum saturation of 204% in the wet season and 160% in the dry season.

**Table 2:** N_2_O concentrations (saturation %) and water-air N_2_O fluxes, for the Johnstone River, the Fitzroy River, and Constant Creek, including area-weighted values.

| Location | $\text{N}_2\text{O}$ saturation (%) | | | | $\text{N}_2\text{O}$ flux ($\mu\text{mol m}^{-2} \text{ day}^{-1}$) | | | |
| --- | --- | --- | --- | --- | --- | --- | --- | --- |
|  | min | max | mean | Area-weighted mean | min | max | mean | Area-weighted mean |
| Johnstone River wet season | 132 | 245 | 170 | **165** | 0 | 47.6 | 6.1 | **5.6** |
| Johnstone River dry season | 97 | 228 | 133 | **140** | $-0.3$ | 28.8 | 2.8 | **3.6** |
| Constant Creek wet season | 93 | 104 | 99 | **100** | $-2.2$ | 1.2 | $-0.1$ | **0.02** |
| Constant Creek dry season | 97 | 132 | 106 | **116** | $-0.8$ | 15.1 | 1.3 | **3.5** |
| Fitzroy River wet season | 100 | 139 | 113 | **107** | $-0.01$ | 33.4 | 3.8 | **2.2** |
| Fitzroy River dry season | 103 | 204 | 128 | **112** | $-4.8$ | 28.5 | 2.6 | **1.5** |

**Figure 3:**
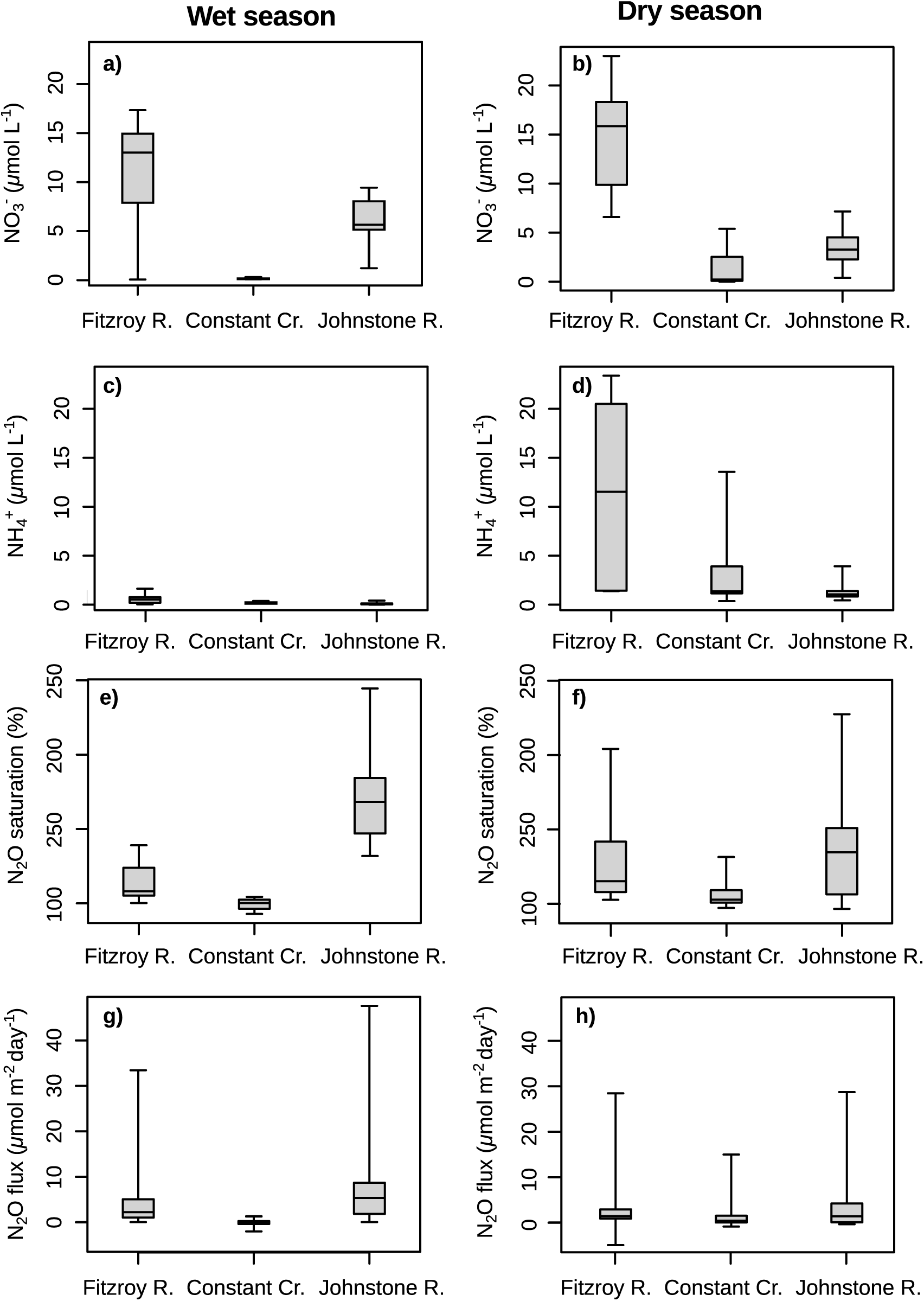
Box plots of wet and dry season NO_3_ ^−^, NH_4_^+^ concentrations, N_2_O saturations, and water-air N_2_O fluxes for the three estuaries.

**Figure 4:**
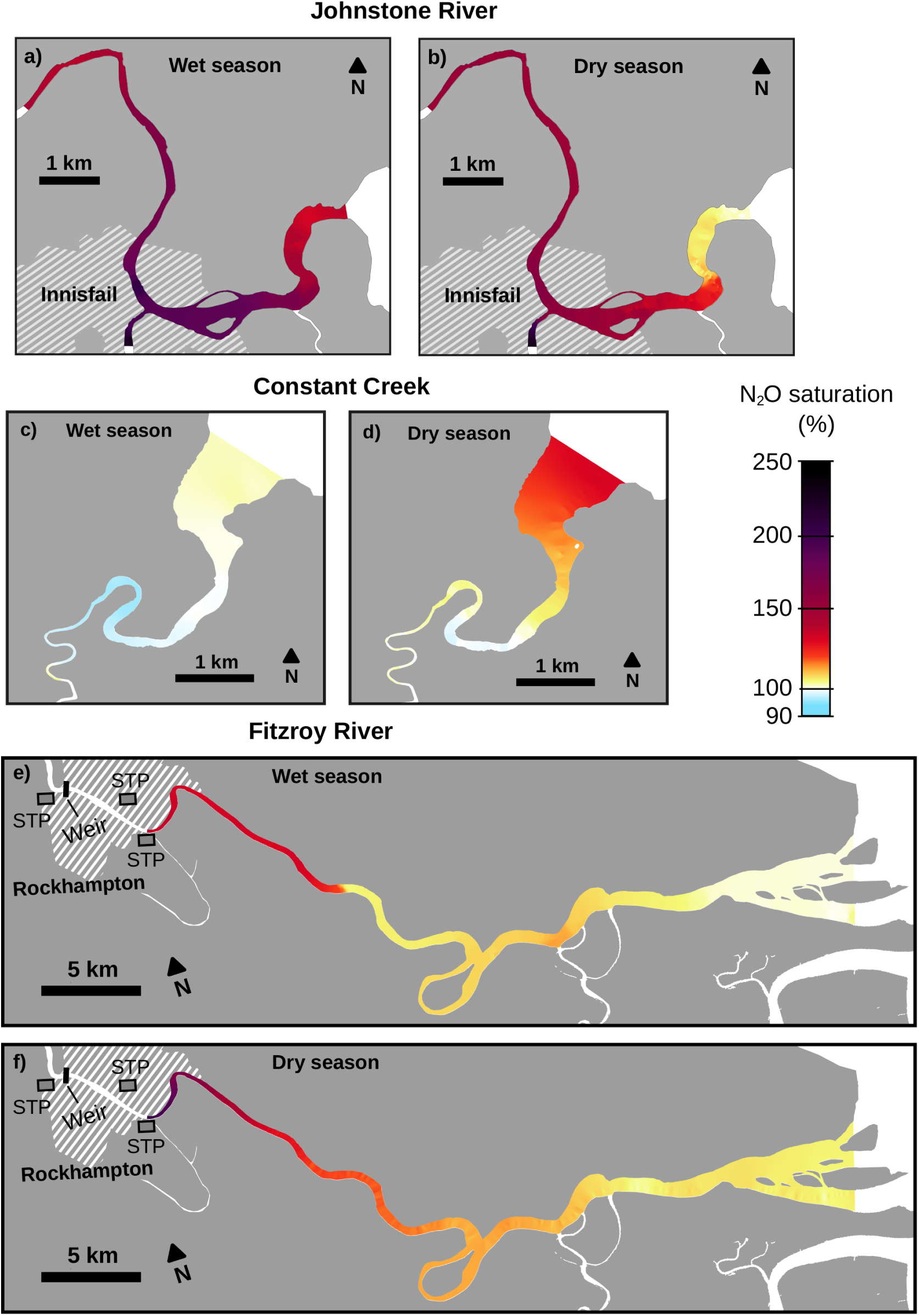
The spatial variability of N_2_O concentration (% saturation) in the Johnstone River Estuary, Constant Creek, and Fitzroy River Estuary in the wet and dry seasons.

The freshwater flushing time in the Johnstone River was 0.59 days (14 h) in the wet season, during which time the theoretical water-column ventilation time was about 2.02 days. The potential contribution of riverine N_2_O evasion to N_2_O emissions was about 450% (Figure 5) corresponding to a total riverine input of 70 mol of excess N_2_O per day, as compared with the total water-air evasion of 15.4 mol d^−1^. In the dry season, the freshwater residence time was 2.3 days, similar to the theoretical ventilation time of 2.26 days. The contribution of riverine N_2_O was 124%; 9.8 mol d^−1^ N_2_O was emitted to the atmosphere, while the total excess freshwater N_2_O contribution was 12.3 mol d^−1^.

**Figure 5:**
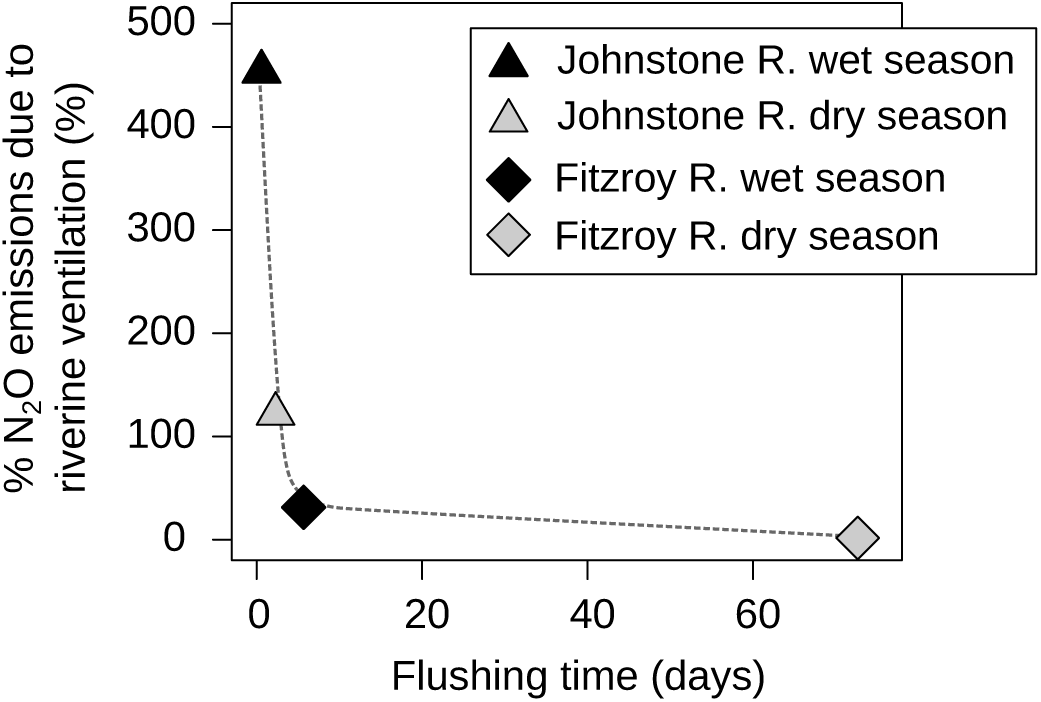
Residence time vs. the proportion of water-air N_2_O flux explained by ventilation of river-derived (freshwater-derived) N_2_O from upstream sources (%), for the Johnstone River Estuary and Fitzroy River Estuary

The Johnstone River was consistently the least saline of the three estuaries, with salinity range of < 1 to 34 in the dry season and < 1 to 12 in the wet season. In both seasons there was a negative relationship between salinity and N_2_O. In both seasons, N_2_O concentrations measured downstream of the confluence of the North and South Johnstone fell below the salinity-N_2_O conservative mixing line, resulting in a total deficit of 6.3 mol in the wet season and 1.4 mol in the dry season (Figure 7). Over the same area, the total water-air N_2_O flux was 8.9 mol d^−1^ (wet season) and 4.2 mol d^−1^ (dry season). Additionally, the freshwater residence times were lower below the confluence of the North and South Johnstone Rivers — 7 h for the wet season and 21 h for the dry season — and over the residence time the water-air N_2_O flux could account for 41% and 260% of the missing N_2_O in the wet and dry season, respectively. The ratio of the N_2_O emissions (*µ*mol d^−1^) to total DIN load (*µ*mol d^−1^) was 0.02% in the wet season and 0.09% in the dry season.

**Figure 6:**
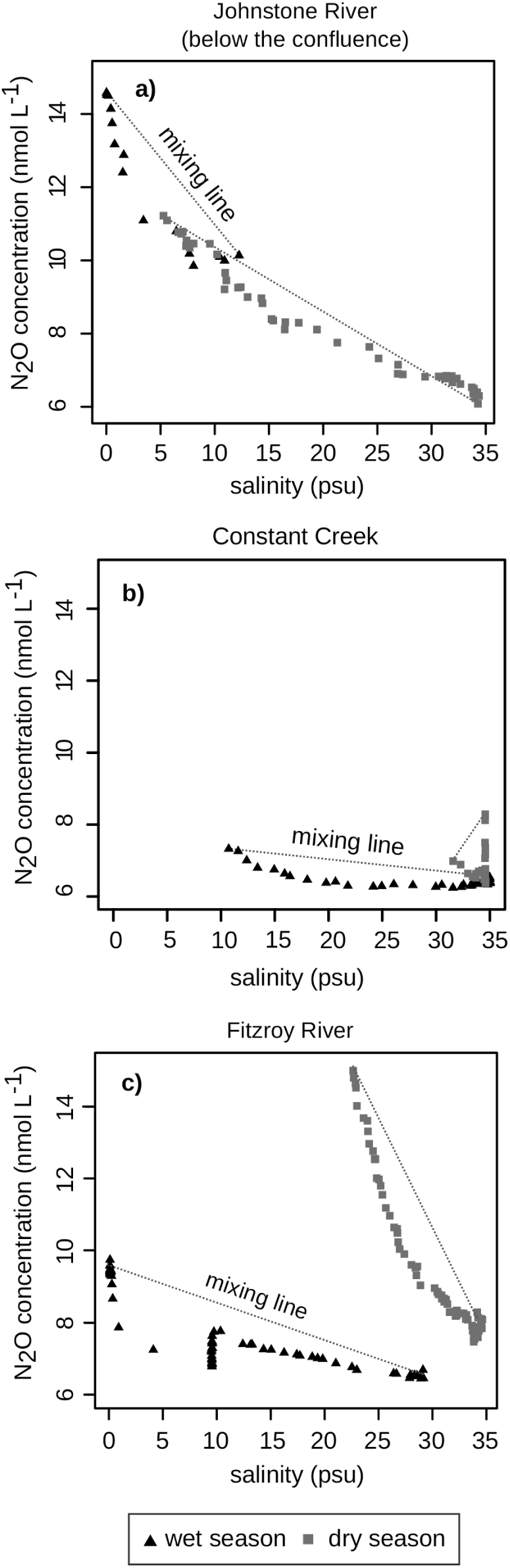
N_2_O concentrations plotted as a function of salinity in wet and dry seasons in the Johnstone River Estuary (a), Constant Creek (b) and Fitzroy River Estuary (c).

**Figure 7:**
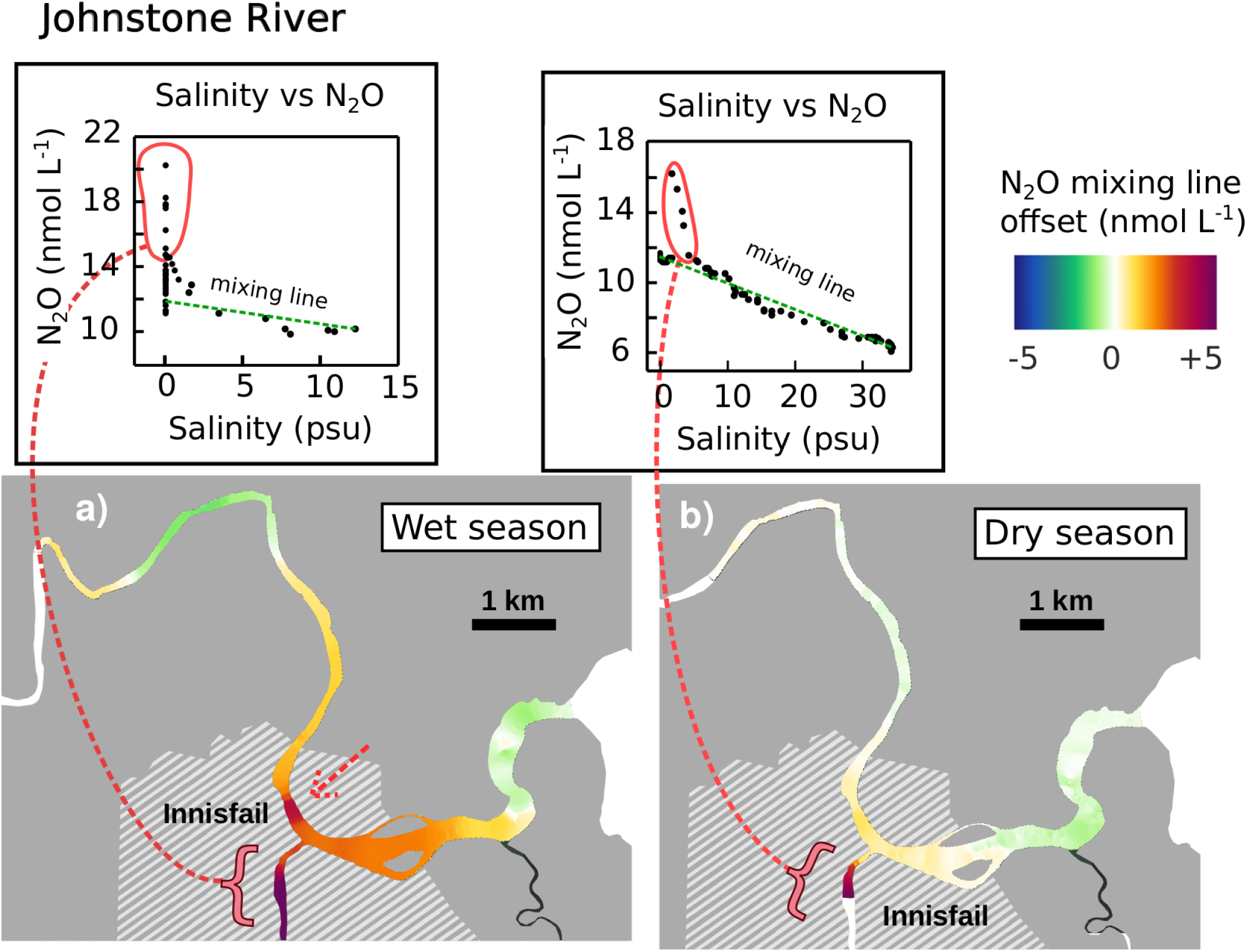
N_2_O mixing plot for the Johnstone River. The map shows the difference, in nmol N_2_O between the measured N_2_O concentration and the value expected from saltwater/freshwater mixing alone, with ‘hot spots’ indicated in red.

In the dry season, NO_3_ ^−^ concentrations were lower (mean value of 3.5 *µ*mol L^−1^) and NH_4_^+^ values higher (mean of 1.3 *µ*mol L^−1^) (Figure 3b,d) than in the wet season (mean of 6.0 *µ*mol L^−1^ for NO_3_ ^−^ and 0.1 *µ*mol L^−1^ for NH_4_^+^) (Figure 3a,c). There was a strong positive correlation between NO_3_ ^−^ and N_2_O in dry season (r^2^ = 0.77; p < 0.001) and a weak and non-significant relationship in the wet season (Figure 8a). Ammonium concentrations were low and showed no correlation with N_2_O concentrations (Figure 8b). Nitrate levels decreased with increasing salinity in both seasons (Appendix Figure A1a), but NH_4_^+^ showed no clear correlation with salinity (Appendix Figure A1b). Dissolved oxygen levels varied between 90% and 100% saturation, with AOU values falling between 7 and 36 *µ*mol L^−1^ in the wet season and – 30 and 16 *µ*mol L^−1^ in the dry season (Appendix Figure A2a,b). Excess N_2_O (ΔN_2_O) ranged from 2 to 10 nmol L^−1^ in the wet season and 0 to 4 nmol L^−1^ L in the dry season.

**Figure 8:**
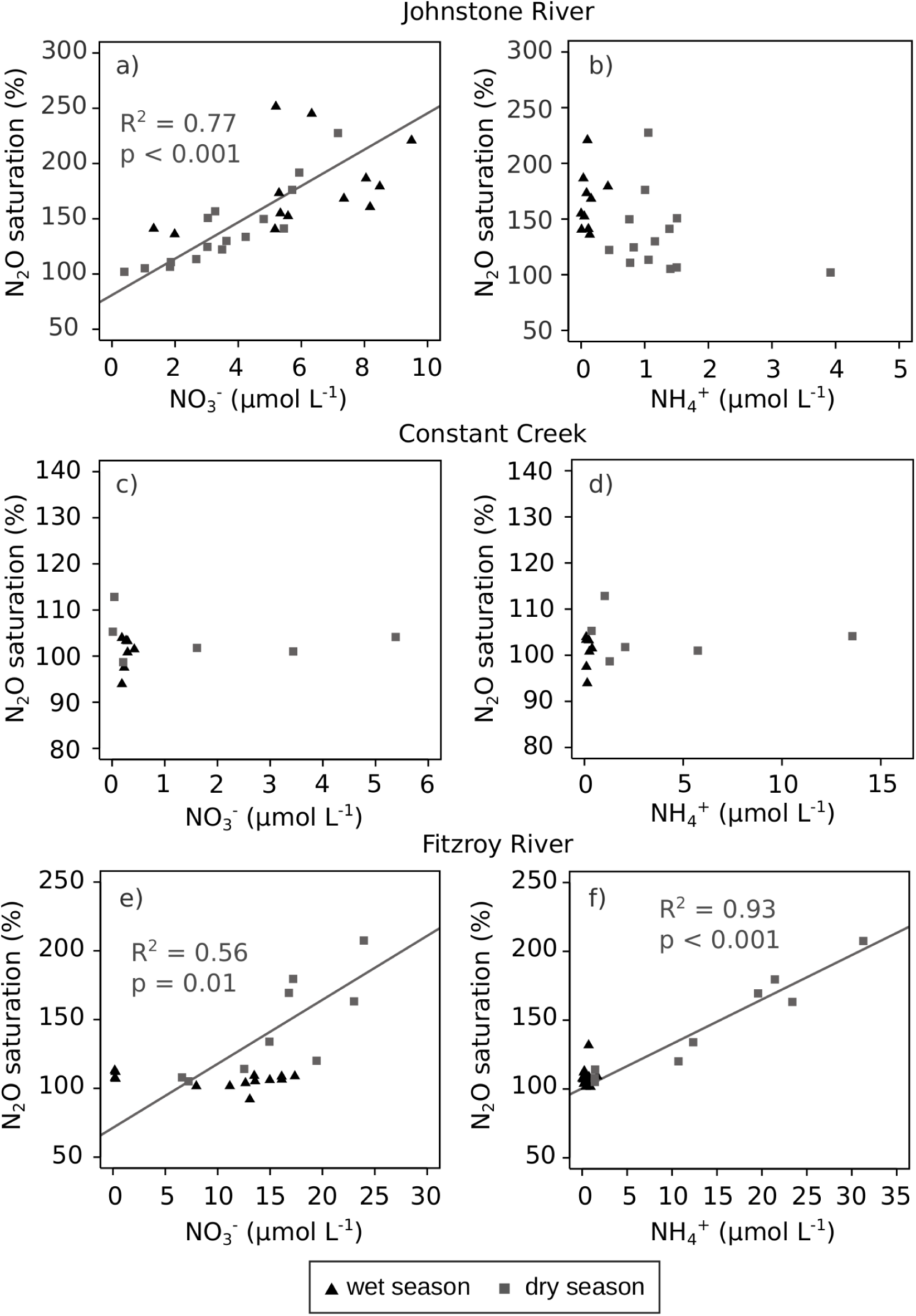
NO_3_ ^−^ and NH_4_^+^ vs N2O concentrations for the Johnstone River Estuary, Constant Creek, and the Fitzroy River Estuary.

The ^222^Rn concentrations were negatively correlated with salinity and positively correlated with both NO_3_ ^−^ and N_2_O concentration in the dry season (Figure 9e,g,h). Overall, wet season *δ* ^15^N-NO_3_ ^−^ values (+ 7.7 to + 9.9‰) were higher than dry season values (+ 4.8 to + 7.3‰) (Figure 10a,b), with the lowest wet season value near the mouth of the estuary, and the highest value in the high-N_2_O reaches of the north arm.

**Figure 9:**
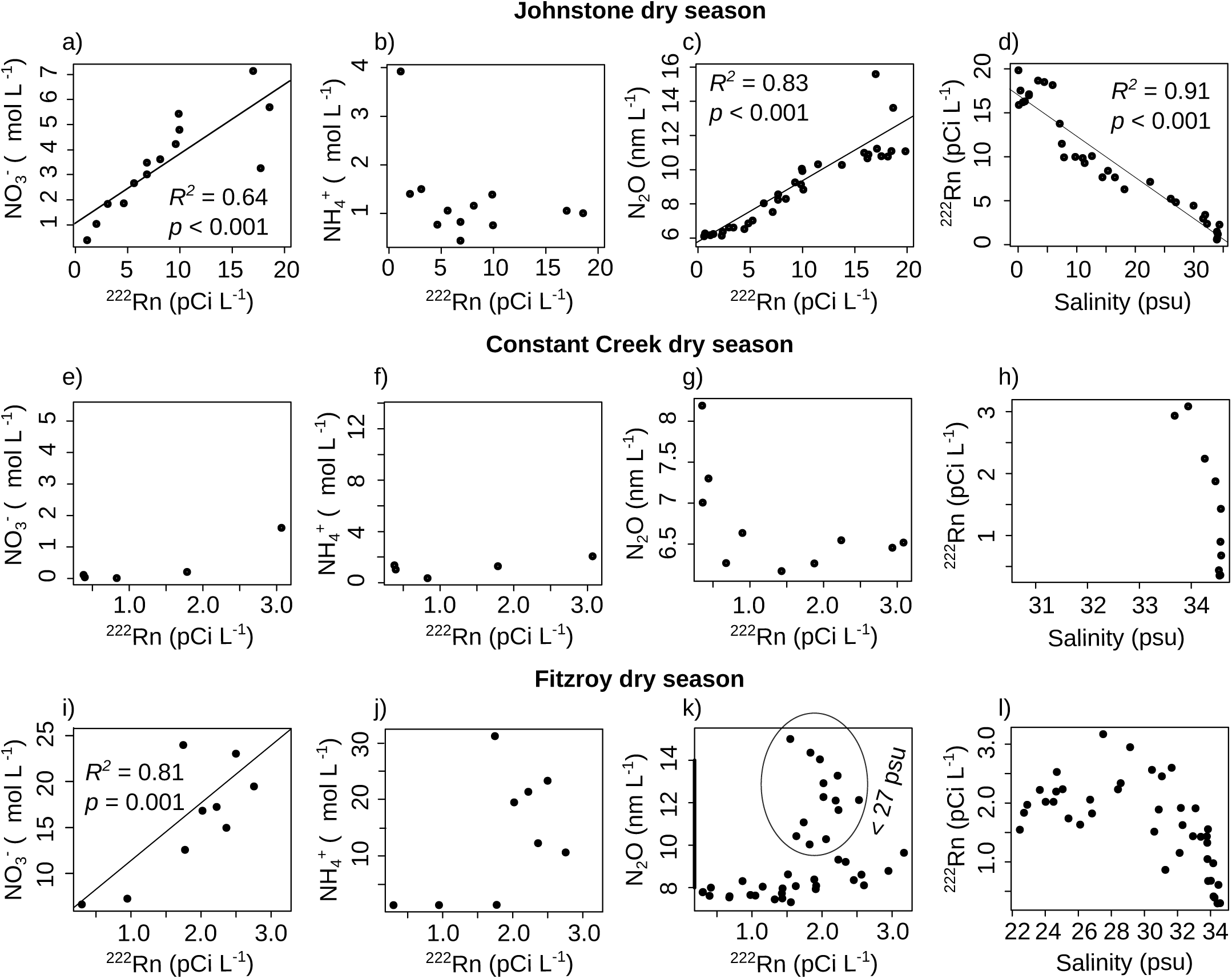
^222^Rn vs. NO_3_ ^−^, NH_4_^+^, and N_2_O (nmol L^−1^) concentrations, and salinity vs. ^222^Rn concentration, for the Johnstone River Estuary, Constant Creek, and the Fitzroy River Estuary

**Figure 10:**
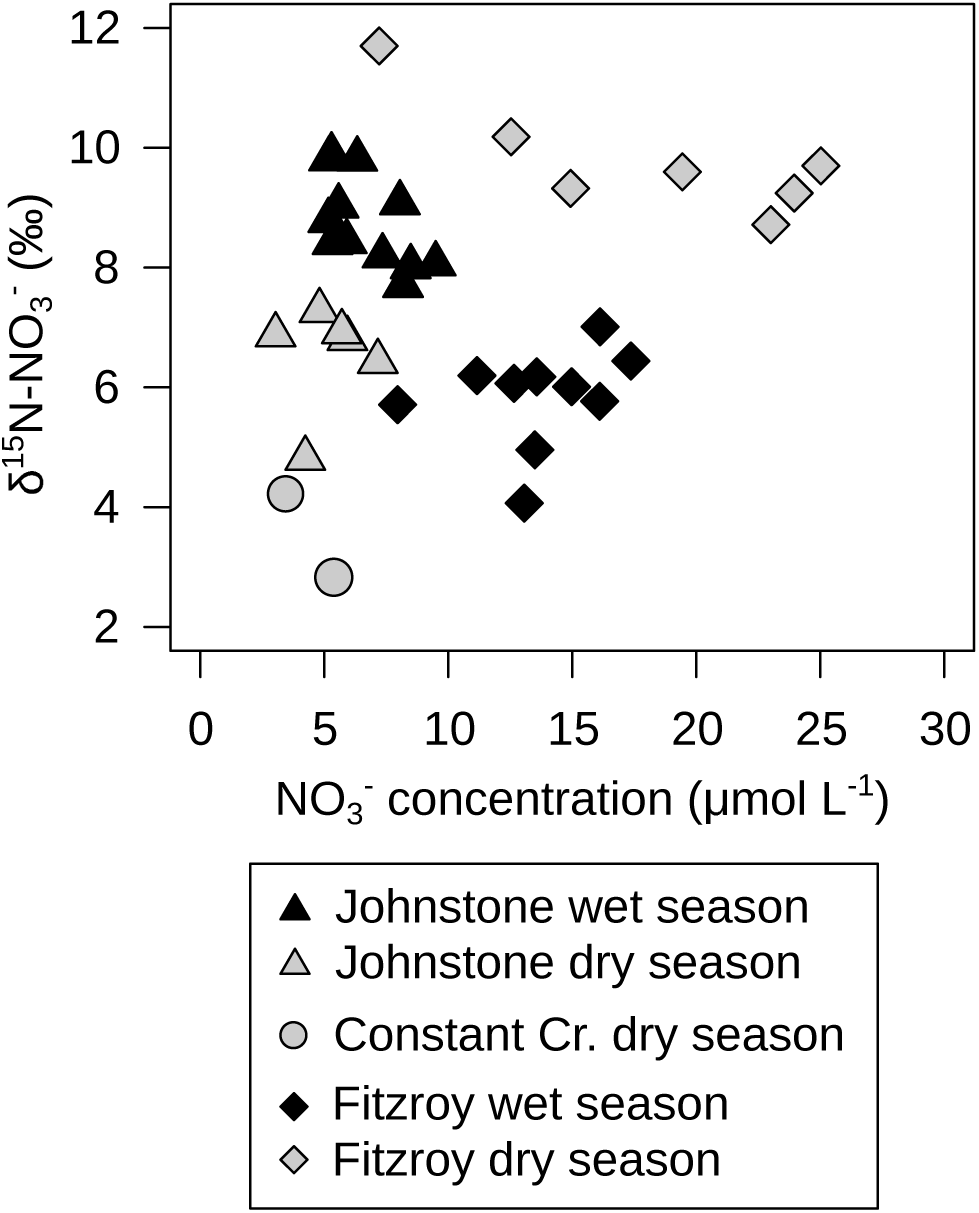
NO_3_ ^−^ concentration vs *δ* ^15^N values of nitrate (*δ* ^15^N −NO_3_ ^−^) during the wet and dry seasons in the Johnstone River Estuary, the Fitzroy River Estuary, and Constant Creek.

### 4.2. Constant Creek Estuary

N_2_O concentrations were lower in Constant Creek than the Johnstone and Fitzroy river estuaries, ranging from 92.9% to 104.4% saturation in the wet season and 97.3% to 131.5% saturation in the dry season (Table 2; Figure 4c,d).

In the dry season N_2_O concentrations were highest at the seawater end-member and showed non-conservative behaviour with an uptake in the middle reaches of the estuary (Figure 4c). In the wet season water-air N_2_O fluxes were negative over a large portion of the estuary (range of – 2.2 to 1.2 *µ*mol m^−2^ d^−1^) (Table 2), reflecting slightly undersaturated N_2_O concentrations (93%-100%) in the middle of the estuary. The highest water-air fluxes (up to 15.1 *µ*mol m^−2^ d^−1^) (Table 2), occurred during the dry season, when N_2_O saturations reached ∼ 130% in the outer estuary (Figure 4d).

Theoretical water-column ventilation times were 2.1 days in the wet season and 1.2 days in the dry season, the difference caused by an increase in wind speed in the dry season. The total water-air flux was much lower in the wet season (1.4 mol d^−1^) than in the dry season (8.9 mol d^−1^). In both seasons, N_2_O concentrations fell below the conservative mixing line, with an apparent total N_2_O deficit of 1.4 mol in the wet season and 9.9 mol in the dry season. For water-air N_2_O emissions to make up for this deficit, it would require ∼ 28 days in the wet season and only ∼ 1 day in the dry season.

Constant Creek was the most saline of the three estuaries, with salinity ranging from 11 to 35 in the wet season and 30 to 35 in the dry season. NO_3_ ^−^ concentrations were lower than in the Johnstone and Fitzroy river estuaries, ranging from 0.2 to 0.4 *µ*mol L^−1^ in the wet season and from 0 to 5.4 *µ*mol L^−1^ in the dry season (Figure 3a,b). Similar to the Fitzroy and Johnstone river estuaries, NH_4_^+^ concentrations were extremely low in the wet season, ranging from 0.1 to 0.4 *µ*mol L^−1^ (Figure 3c), but increased in the dry season, ranging from 0.4 to 13.6 *µ*mol L^−1^ (Figure 3d). In the wet season the highest NO_3_ ^−^ and NH_4_^+^ concentrations were at the seawater end-member. In contrast, in the dry season the highest NO_3_ ^−^ and NH_4_^+^ concentrations were at the river end-member (Appendix Figure A1c,d).

Neither N_2_O, NO_3_ ^−^ nor NH_4_^+^ were correlated with ^222^Rn (Figure 9i,j,k); ^222^Rn increased at lower salinities (Figure 9l) while DIN and N_2_O concentrations were highest in the outer estuary (Figure 4d). In the dry season dissolved oxygen was saturated (∼ 100%), with AOU values between – 8 and 13 *µ*mol L^−1^ (Appendix Figure A2d). In the wet season dissolved oxygen was slightly undersaturated (85%), with AOU values between – 19 and 34 *µ*mol L^−1^. AOU and ΔN_2_O show a negative correlation in the wet season (Appendix Figure A2c). Due to low NO_3_ ^−^ concentrations, *δ* ^15^N-NO_3_ ^−^ values were only measured at two locations during the dry season, when they were 2.8 and 4.2‰.

### 4.3. Fitzroy River Estuary

In the Fitzroy River Estuary, N_2_O concentrations ranged from 100% to 139% saturation, in the wet season and 103% to 204% in the dry season (Table 2; Figure 3e,f). Despite higher N_2_O concentrations in the dry season, water-air fluxes were higher in the wet season (area-weighted mean of 2.2 *µ*mol m^−2^ d^−1^ in wet and 1.5 *µ*mol m^−2^ d^−1^ in dry) due to higher k-values during the wet season (Appendix Table A2). In both seasons N_2_O concentrations decreased from the upstream end to the ocean. However in the wet season there was a slight increase in the middle reaches of the estuary, near the mouth of small mangrove-lined tributary at about 30 km downstream of Rockhampton.

The freshwater flushing time was 5.7 days in the wet season, and the average ventilation time of the water-column was 0.9 days. With a freshwater excess N_2_O load of 151 mol N_2_O and a total water-air N_2_O flux of 477 mol over the residence time, the ventilation of the freshwater (riverine) N_2_O accounted for 32% of the N_2_O emissions (Figure 5). In the dry season the flushing time was 72.6 days and the ventilation time was 2.2 days. The excess freshwater N_2_O load was 67 mol, and over the duration of the residence time, the estimated total water-air flux was 4150 mol, meaning that ventilation of river N_2_O inputs can explain 1.6% of the water-air flux. In both seasons, N_2_O concentrations showed non-conservative mixing, (Figure 6c); N_2_O was as much as 3 nmol L^−1^ below the conservative mixing line despite N_2_O saturation remaining above 100%. The estimated total N_2_O deficit was 1.3 mol N_2_O in the wet season and 150 mol in the dry season. Over the duration of the residence time, the water-air flux could account for 5100% and 2800% of the lost N_2_O in the wet season and dry season, respectively. The N_2_O water-air flux (*µ*mol d^−1^) accounted for 0.05% and 1.12% of the total DIN load (*µ*mol d^−1^) in the wet and dry seasons respectively.

Salinity was lower during the wet season, ranging from < 1 to 29 as compared to 24 to 35 in the dry season. In the dry season, NO_3_ ^−^ and NH_4_^+^ concentrations were elevated (Figure 3b,d) compared to the Johnstone River Estuary and Constant Creek, showing a positive correlation with salinity (Appendix Figure A1e,f). The N_2_O saturation was positively correlated with both NO_3_ ^−^ and NH_4_^+^ concentrations in the dry season (Figure 8e,f). In contrast, in the wet season NO_3_ ^−^ and NH_4_^+^ concentrations were lower (Figure 3a,c) and not correlated with N_2_O concentrations (Figure 8e,f).

AOU values varied from 24 to 59 *µ*mol L^−1^ in the wet season, increasing along the length of the freshwater plug until it reached a peak at the transition between freshwater (salinity < 0.5) and brackish water (Appendix Figure A2e). AOU increased again until about 30 km from Rockhampton, where both AOU and ΔN_2_O reach a secondary, mid-estuary peak. In the dry season, AOU fell between 14 and 41 *µ*mol L^−1^. The highest AOU values were observed at a salinity less than 25, where ΔN_2_O was elevated in the upper estuary. At this location there was a positive correlation between AOU and salinity (Appendix Figure A2f). At a salinity of ∼ 27 and a distance of ∼ 11 km downstream of Rockhampton, the AOU–ΔN_2_O relationship changes, but the values further downstream still show a positive correlation.

The ^222^Rn concentrations showed a positive correlation with NO_3_ ^−^ concentrations (Figure 9i) and a maximum at intermediate salinity values — (up to 3 pCi L^−1^ between salinities of 26 and 28) (Figure 9l). At the lowest salinity values (< 25), where ^222^Rn was at intermediate concentration, NO_3_ ^−^, N_2_O and NH_4_^+^ were all relatively high, diverging from a simple linear correlation with ^222^Rn. Nitrate *δ* ^15^N values were higher in the dry season (+ 8.7 to + 11.7‰), than in the wet season (+ 4.1 to + 7.0‰) (Figure 10) and the lowest wet-season *δ* ^15^N-NO_3_ ^−^ values were found in the low-salinity upper reaches of the estuary.

## 5. Discussion

### 5.1. Johnstone River Estuary

The highest concentrations of N_2_O (both wet and dry seasons) over the three estuarine systems were found in the Johnstone River Estuary. In contrast to the two other systems, N_2_O concentrations in the Johnstone River Estuary were higher during the wet season rather than the dry season. Below we discuss these differences in terms of the potential sources of N_2_O, the observed seasonal pattern in N_2_O concentration, and the distribution of N_2_O within the Johnstone River Estuary.

Compared with the other two estuaries, the Johnstone River Estuary is subject to much higher freshwater input from its tributaries (Figure 2a,b,c). As such, the estimated N_2_O loads are high enough, and the residence time is low enough, that some of the N_2_O arriving from upstream sources is exported to the ocean. In both the wet and dry season, the proportion of N_2_O emissions explained by ventilation of riverine N_2_O inputs is greater than 100% (Figure 5). In other words the potential N_2_O ventilation is higher than the observed N_2_O water-air flux, and any in situ production of N_2_O is overwhelmed by the river load.

Discharge-driven increases in estuarine N_2_O concentration are relatively common. For example, across 28 estuaries in India, high wet-season N_2_O concentrations resulted from mobilisation of N_2_O from upstream sources (Rao and Sarma, 2013). Similar to the Johnstone River, the Saribas and Lupar Rivers in Borneo also exhibited greater N_2_O variability and higher maximum N_2_O concentrations in the wet season, with a clear source of N_2_O near the midpoint of the spatial transect (Mueller et al., 2016). In the dry season however, Mueller et al. (2016) found that N_2_O concentrations were less variable and most of the N_2_O came from upstream, freshwater inputs.

One of the main differences between the Johnstone River and the two rivers in Borneo is the relationship between N_2_O and NO_3_ ^−^. Mueller et al. (2016) found no correlation between DIN and N_2_O in either season, and noted that despite the relatively low DIN concentrations of the waterways (10–37 *µ*mol L^−1^) N_2_O was present in very high concentrations in some places (up to 1679% saturation). In contrast, in the Johnstone, N_2_O concentrations showed a correlation with NO_3_ ^−^ in both seasons (Figure 8a,b), indicating that N_2_O production was enhanced by the higher NO_3_ ^−^ concentrations, or that N_2_O and NO_3_ ^−^ were entering the system together.

The salinity-NO_3_ ^−^ mixing plots indicate that most of the NO_3_ ^−^ in the Johnstone River was from freshwater sources from the North and South Johnstone tributaries (Appendix Figure A1a); NO_3_ ^−^ was not generated in large quantities below the confluence of the river. It is difficult to tell if any N_2_O was produced in the estuary. In both seasons, the N_2_O load from freshwater input is high enough to more than account for all of the emitted N_2_O (Figure 5) and in-situ production is not needed to explain the observed patterns. In some estuaries and coastal waters, a positive correlation between N_2_O and apparent oxygen utilization has been interpreted as a sign of in-situ nitrification (Nevison et al., 2003; Goncalves and Brogueira, 2017), however this correlation was not observed in the Johnstone (Figure A2a,b), and neither did we observe a negative correlation between N_2_O and NH_4_^+^ (Figure 8b), as would be expected if nitrification were consuming large quantities of NH_4_^+^.

Another possible pathway of N_2_O production or consumption is denitrification. Sediments and groundwater can be major sites of denitrification and N_2_O production (Marzadri et al., 2017), and the higher *δ* ^15^N-NO_3_ ^−^ values observed in the wet season could indicate a greater influence of denitrification during the high-N_2_O season (Figure 10). Denitrification increases the *δ* ^15^N-NO_3_ ^−^ in the remaining NO_3_ ^−^ pool (Granger et al., 2008; Middelburg and Nieuwenhuize, 2001), therefore the higher *δ* ^15^N-NO_3_ ^−^ could result from an increased inflow of DIN from sediment pore-water or groundwater, where denitrification is likely to occur. However, similar to nitrification, there was little evidence that denitrification occurred in-situ; dissolved N_2_O could have come from upstream sources.

A denitrification pathway for N_2_O production would be consistant with the strong correlation between ^222^Rn and both NO_3_ ^−^ and N_2_O in the dry season (Figure 9a,c), which suggest that at least some NO_3_ ^−^ and N_2_O came from groundwater inputs. The CO_2_ and CH_4_ concentrations were also correlated with ^222^Rn suggesting a similar groundwater source (Rosentreter et al., 2018). Although we were unable to obtain ^222^Rn concentrations for the wet season, it seems likely that groundwater input would increase with higher precipitation. This would drive pore-water NO_3_ ^−^ and N_2_O into the upper river systems. Overall, we propose that the higher concentration of N_2_O in the Johnstone River Estuary relative to the Constant Creek and the Fitzroy River Estuaries, and the increase in N_2_O concentration in the wet season relative to the dry season, is a consequence of higher rainfall and increased groundwater flow.

#### 5.1.1. Spatial Variability

The Johnstone Estuary itself has two main sections (see Figure 1): the North and South tributaries. The South Johnstone tributary was the strongest source of N_2_O-rich freshwater in both the wet and dry seasons, accounting for 51–55% of the N_2_O load despite only contributing 32–34% of the freshwater volume (according to the upstream gauges). This high N_2_O concentration water mixed within the main body of the estuary with water from the North Johnstone tributary.

Below the confluence of the North and South Johnstone, N_2_O concentrations fall below the conservative mixing line in both the wet and dry seasons (Figure 6a). If N_2_O concentrations were near or below 100% saturation, this mid-estuary minimum of N_2_O might indicate N_2_O reduction to N_2_, as has been observed in some low-DIN tidal waterways (Daniel et al., 2013; Rao and Sarma, 2013; Erler et al., 2015; Maher et al., 2016; Murray et al., 2018). However, where N_2_O is supersaturated, as in the Johnstone River Estuary, it doesn’t seem likely that N_2_O consumption could explain the loss of N_2_O.

The N_2_O-salinity relationship in the Johnstone River Estuary is similar to the eutrophic Pearl River in China (Lin et al., 2016), where N_2_O was non-conservative despite dissolved N_2_O concentrations consistently above 100% saturation (101–3799%). Lin et al. (2016) estimated N_2_O removal rates which nearly balanced out water-air N_2_O fluxes, suggesting that all of the apparent N_2_O loss could be explained by evasion.

In the Johnstone River, in contrast, there was a net export of freshwater-derived N_2_O (55 mol d^−1^) in the wet season, despite the fact that evasion could only explain ∼ 40% of the N_2_O lost below the confluence of the North and South Johnstone. In the dry season, this N_2_O loss was dwarfed by the water-air evasion (280%), a fact that might indicate in-situ N_2_O production, yet the net export of freshwater-derived N_2_O was lower (2.4 mol d^−1^). The residence time was very low below the confluence of the North and South Johnstone in both seasons, and a small change in either residence time or water volume could have a large effect on the calculations.

### 5.2. Constant Creek Estuary

In contrast to the Johnstone River, the low DIN and N_2_O concentrations in Constant Creek suggest relatively low anthropogenic input of nitrogen from agriculture or other sources. Similar to the Johnstone River Estuary, N_2_O concentrations in Constant Creek fall below the conservative mixing line (Figure 6c). In this case, however, N_2_O concentrations in the middle reaches of the estuary are close to or slightly below 100% saturation. This suggests that the N_2_O deficit results from consumption of N_2_O by denitrifiers.

In the wet season there was some evidence for in-situ consumption. While the overall area-weighted flux was positive, negative water-air fluxes (as low as – 2.2 *µ*mol m^−2^ d^−1^) were observed in the middle section of the transect (Table 2). The undersaturation of N_2_O has been observed previously in other relatively pristine creeks in Australia (Erler et al., 2015; Maher et al., 2016; Murray et al., 2018; Wells et al., 2018), a salt marsh (Daniel et al., 2013), and some monsoonal estuaries in India (Rao and Sarma, 2013). However, N_2_O undersaturation is not observed in all pristine estuaries. Barnes et al. (2006) documented N_2_O saturations between 100% and 208% in a pristine mangrove creek where NO_3_ ^−^ values were 0–5 *µ*mol L^−1^ and NH_4_^+^ values were 0–25 *µ*mol L^−1^, similar to values found at Constant Creek (0–5.4 *µ*mol L^−1^ NO_3_ ^−^ and 0–13.6 *µ*mol L^−1^ NH_4_^+^). The creek on Andaman Island had relatively low salinity (0–28) compared with Constant Creek (10–35); the shared characteristics of undersaturated estuaries seem to be low DIN and high salinity (Maher et al., 2016; Rao and Sarma, 2013). Saline conditions can inhibit water-column nitrification (Rysgaard et al., 1999), and this might help to prevent in-situ N_2_O production.

In the dry season, it was not clear that N_2_O was consumed in the estuary, due to the fact that that N_2_O concentrations were generally elevated above 100% (Figure 3f). Constant Creek was the only estuary where the highest N_2_O concentrations were found at the downstream end of the transect, and we propose that the extra N_2_O was most likely produced on the extensive shallow shoals which are found in the outer estuary (Figure 4d). Shallow benthic sediments are known to be a strong source of N_2_O in many estuaries (e.g. Usui et al., 2001; Ferrón et al., 2007). Due to the downstream N_2_O source, the salinity-N_2_O data-points to deviate below the conservative mixing line. Considering that N_2_O concentrations remain at or above 100% over the transect, we cannot say that the non-conservative behaviour of N_2_O is due to in-situ consumption. Furthermore, the water-air N_2_O flux would require only 1.1 days to balance out the negative deficit, which is not an unreasonably long time.

In the dry season, ^222^Rn concentrations were highest at low N_2_O concentrations, indicating that groundwater was not a strong source of N_2_O (Figure 9g). Water-column nitrification is also unlikely to produce N_2_O in this environment. As mentioned previously, a positive linear correlation between AOU and ΔN_2_O is considered a sign of nitrification activity because nitrification produces N_2_O and consumes O_2_. However in Constant Creek there is a clear negative correlation between excess N_2_O and AOU in the wet season (Appendix Figure A2), suggesting that nitrification was not a strong source of N_2_O, but that denitrifiers reduced N_2_O to N_2_, especially under low-DIN conditions.

### 5.3. Fitzroy River Estuary

In the Fitzroy River Estuary, there was a clear source of N_2_O at the upstream end of the survey (Figure 4e,f), with higher concentrations during the dry season (Figure 4f). Similar to Constant Creek, but in contrast to the Johnstone River Estuary, the seasonal differences in N_2_O concentrations reflect a fundamental shift in the source and/or processing of DIN, and not just a change in freshwater discharge. While total freshwater discharge was higher in the wet season, this extra freshwater did not have enough N_2_O to increase N_2_O concentration higher than in the dry season (Figure 6c), and freshwater may even dilute N_2_O concentrations. In the dry season the weak relationship between N_2_O and ^222^Rn shows that pore-water/groundwater was not the only source of N_2_O in the Fitzroy River Estuary (Figure 9k). Near Rockhampton, there was a source of low-salinity water with relatively low ^222^Rn concentrations but high N_2_O, NO_3_ ^−^ and NH_4_^+^ concentrations (Figure 9i,j,k,l).

Looking at the strong correlation between N_2_O saturation and DIN (Figure 8e,f) and the relationship between ^222^Rn and DIN, it seems likely that the wastewater treatment plants located near the head of the Fitzroy River Estuary Figure 1) are the ultimate source of a large portion of the NO_3_ ^−^, N_2_O and NH_4_^+^ present in the waterway. In the case of N_2_O, the treatment plants may be a direct or indirect source: N_2_O may be discharged directly as a result of wastewater processing, or alternately the organic matter and DIN from the wastewater treatment plants may spur N_2_O production downstream of the wastewater outlet. The low-salinity, low-^222^Rn wastewater had a greater effect on NH_4_^+^ and N_2_O concentrations than on NO_3_ ^−^ concentrations (Figure 9i,j,k). For a given NO_3_ ^−^ concentration in the dry season N_2_O concentrations were much lower in the Fitzroy than in the Johnstone where NO_3_ ^−^ and N_2_O were sourced from groundwater (Figure 3b,f). This suggests that although most of the N_2_O and some of the NO_3_ ^−^ were sourced from the wastewater treatment plant the N_2_O was not directly linked to the NO_3_ ^−^. Nitrification of wastewater-derived NH_4_^+^, occurring at or just downstream of the wastewater outlet could also have elevated N_2_O and NO_3_ ^−^.

Wastewater effluent is associated with increased NH_4_^+^ concentrations and N_2_O production in other tropical and subtropical estuaries. For example, in the Adyar River in southeast India high NH_4_^+^ concentrations resulted from wastewater and use of diammonium phosphate fertiliser led to elevated N_2_O driven by water-column nitrification (Rajkumar et al., 2008). Many of the estuaries on the east and west coast of India, where N_2_O production is attributed to water-column nitrification, receive wastewater effluent as well and show a strong correlation between NH_4_^+^ and N_2_O concentration (Rao and Sarma, 2013). Similarly, strong DIN-N_2_O correlations were observed in the wastewater-affected Pearl River in China (Lin et al., 2016).

Upstream freshwater N_2_O inputs could not account for the entire water-air N_2_O flux in either season, which indicates that there is a secondary source of N_2_O downstream of Rockhampton, either from tributary/groundwater inputs or in-situ production. In the wet season, some of this extra N_2_O entered the estuary about 30 km downstream from Rockhampton, where there was an obvious increase in N_2_O concentration over a region of constant salinity. Considering that this occurs near the mouth of a creek where we observed high N_2_O concentrations, it seems likely that some of this extra N_2_O was allochthonous. However the N_2_O increase was also associated with an increase in AOU, indicating a possible role of nitrification. AOU declines again downstream of the mid-estuary N_2_O peak, indicating that nitrification was likely responsible for very little N_2_O production in the low-N_2_O lower reaches of the estuary.

In the dry season, the total contribution of freshwater-derived N_2_O is not well known because freshwater N_2_O concentrations were not directly measured (minimum salinity in the estuary is 22). Additionally, estimating ‘freshwater’ N_2_O concentration is difficult where wastewater inputs contribute to N_2_O production, as wastewater could have an extremely high N_2_O concentration. While excess N_2_O (ΔN_2_O) was assumed to be similar to the highest observed values at the upstream end of the estuary (8.0 nmol L^−1^), if we assume the average freshwater ΔN_2_O was close to wet season values (∼ 2.8 nmol L^−1^), the riverine contribution to N_2_O production would be 0.4%. At the high value of 20 nmol L^−1^ the contribution would be 4%, and to reach 100%, the average freshwater ΔN_2_O would have to be ∼ 500 nmol L^−1^.

Unlike the wet season, the mid-estuary tributaries don’t seem to have a huge effect on N_2_O concentration. Above the salinity of ∼ 27 transition at ∼ 11 km from Rockhampton, some N_2_O production could result from nitrification; AOU concentrations were slightly higher at salinities below 25 (up to 1.3 mg L^−1^) where they exhibited a positive relationship with ΔN_2_O (Appendix Figure A2f). However, given that low-O_2_ waters were likely quickly mixed with more oxygenated waters further downstream, it is possible that the AOU−ΔN_2_O pattern is merely the result of mixing and N_2_O evasion, which could cause both AOU and ΔN_2_O to decline.

Considering the estimated freshwater N_2_O inputs and water-air fluxes, it seems that the Fitzroy was the only estuary where residence times were long enough for significant in-situ N_2_O production to occur in both seasons. This raises the possibility that denitrification at the sediment-water interface could also contribute N_2_O. Previous research has identified a relationship between residence time and denitrification in temperate estuaries (Eyre et al., 2016; Nixon et al., 1996), which would predict that over a residence time of 72 days (as in the dry season) about 25% of NO_3_ ^−^ inputs should be consumed by denitrifiers. For residence times of less than 1 month, as in the wet season and in the Johnstone River, the predicted consumption of NO_3_ ^−^ by denitrifiers would be much lower than 25%. In theory, at higher temperatures these values could increase (Barnes and Owens, 1999), however we should note that NO_3_ ^−^ concentrations do not seem to show non-conservative behaviour indicative of significant NO_3_ ^−^ consumption along the length of the estuary (Appendix Figure A1a).

In contrast to the Johnstone River, the *δ* ^15^N−NO_3_ ^−^ data in the Fitzroy seem to indicate more intense denitrification of NO_3_ ^−^ during the dry season, rather than the wet season, which could reflect processing during wastewater treatment and subsequent NO_3_ ^−^ consumption by denitrifiers in the sediment downstream of the wastewater inlet (Figure 10c,d). Without the wastewater source and long residence time, we might expect that the Fitzroy dry season *δ* ^15^N-NO_3_ ^−^ values would be lower than the wet season values, perhaps similar to the dry season values observed at Constant Creek. However the Fitzroy is similar to the other two estuaries in that the seasonal increase in NO_3_ ^−^ input is associated with increased *δ* ^15^N-NO_3_ ^−^ values and higher N_2_O concentrations.

### 5.4. Nitrous oxide load and residence time

Residence time is an important control on the amount of nitrogen and phosphorus processing that occurs in estuaries, with greater processing as residence time increases (Balls, 1994; Nixon et al., 1996; McKee et al., 2000; Eyre et al., 2016). Here, we show that residence time is also an important control on the freshwater N_2_O load that is lost as an estuarine water-air N_2_O flux (i.e. residence time vs. the % riverine ventilation; Figure 5). This is similar to the findings that residence time affects the amount of freshwater DIC that is lost as a water-air CO_2_ flux, (i.e. residence time vs. the % riverine ventilation) (Borges and Abril, 2011).

In the Johnstone River, N_2_O was delivered along with freshwater (surface water and/or groundwater), causing a clear negative relationship between N_2_O concentration and salinity (Figure 6a) and a wet-season dominance of N_2_O (Figure 4a,b). This is typical of many temperate (Barnes and Owens, 1999; Robinson et al., 1998; Zhang et al., 2010) and some tropical estuaries (Rao and Sarma, 2013); in these cases, the dominant control on N_2_O concentration is delivery of freshwater rather than in-situ N_2_O production. In the Johnstone River, the dominance of freshwater-derived N_2_O can be seen in Figure 5, where the amount of freshwater-derived N_2_O exceeds the total water-air N_2_O flux over the surface area of the transect, i.e. the proportion of water-air fluxes that can be explained by riverine N_2_O inputs is greater than 100%.

When residence time increases, however, freshwater inputs play a lesser role compared with in-situ processes. For example, when N_2_O concentrations are higher in the dry season (Chen et al., 2015; Rajkumar et al., 2008; Senthilkumar, 2008), often N_2_O production is attributed to stagnation of DIN-rich water in river or estuary water bodies, leading to water-column nitrification. In the wet season, the in-situ nitrification source of N_2_O is suppressed because of the lower residence time and higher flushing rate (Chen et al., 2015). A similar pattern is evident in the Fitzroy River, and can be seen in the relationship between residence time and % riverine N_2_O ventilation (Figure 5). In the wet season the larger freshwater input dilutes the wastewater effluent and decreases NO_3_ ^−^, NH_4_^+^ and N_2_O concentrations in the estuary. This leads to a lower residence time and a higher proportion of freshwater-derived N_2_O, even though overall N_2_O concentrations are lower (Figure 4c,d). In the dry season, freshwater input is low and the residence time is high (∼ 72 days), strengthening the relative contribution of in-situ production to the water-air N_2_O flux.

### 5.5. Role of water column DIN concentrations and N_2_O emissions

The low Fitzroy River Estuary and Johnstone River Estuary N_2_O concentrations were typical of some tropical or subtropical estuaries or creeks at lower salinity which have a low freshwater source of DIN (Barnes et al., 2006; Maher et al., 2016; Senthilkumar, 2008; Wells et al., 2018) (Table 1). For example, most estuaries surveyed in India exhibited N_2_O saturations between 100% and 300% (Rao and Sarma, 2013), with the exception of a few which had large anthropogenic N inputs. Similar values were observed in the more saline, down-stream reaches of more eutrophic estuaries (Lin et al., 2016; Mueller et al., 2016; Rao and Sarma, 2013), where a strong upstream N_2_O source is diluted by mixing with seawater. More anthropogenically disturbed tropical estuaries, with a large population living in the catchment and significant wastewater or agricultural DIN inputs, can reach much higher maximum saturations, typically at the upstream, freshwater end of the estuary (Chen et al., 2015; Lin et al., 2016; Liu et al., 2015; Rao and Sarma, 2013).

Like N_2_O saturations, calculated N_2_O water-air fluxes across the three estuaries were at the low end of the range typical for estuarine waters. This reflects relatively low DIN concentrations across the three estuaries (below 30 *µ*mol L^−1^ for both NO_3_ ^−^ and NH_4_^+^). Over a large range of NO_3_ ^−^ concentrations (i.e. 0 to 500 *µ*mol L^−1^ of NO_3_ ^−^) water-column NO_3_ ^−^ concentrations appear to be a driver of N_2_O water-air fluxes (Figure 11a) (Murray et al., 2015). Including more recent work in mangrove creeks (Daniel et al., 2013; Erler et al., 2015; Maher et al., 2016; Murray et al., 2018) and other low concentration estuaries (Wells et al., 2018), it appears that negative water-air fluxes occur where water column NO_3_ ^−^ concentration falls below ∼ 5 *µ*mol L^−1^ (Figure 11b), despite the weaker relationship over the smaller NO_3_ ^−^ range. NO_3_ ^−^ concentrations below 5 *µ*mol L^−1^ are commonly found in many undisturbed and slightly disturbed tropical estuaries and mangrove creeks (Table 3), suggesting N_2_O uptake may be widespread.

**Table 3:** NO_3_^−^ and NH_4_^+^ concentrations in tropical marine systems.

| Citation | Environment | Country | $\text{NH}_4^+$ ($\mu\text{mol L}^{-1}$) | $\text{NO}_3^-$ ($\mu\text{mol L}^{-1}$) |
| --- | --- | --- | --- | --- |
| Agawin et al. (1996) | seagrass | Phillippines | 1.6 to 1.9 | 0.4 to 0.8 |
| Carruthers et al. (2005) | seagrass | Panama | 0.20 to 0.26 | 0.20 to 0.27 |
| Erfteimeijer and Herman (1994) | seagrass | Indonesia | 2.2 to 3.2 | 0.9 to 1.4 |
| Heminga et al. (1994) | seagrass | Kenya | 0.4 to 0.8 | 0.3 to 0.45 |
| Uku and Björk (2005) | seagrass |  | 1.6 to 1.9 | 0.4 to 0.8 |
| Boto and Wellington (1988) | mangrove | Australia | 0.1 to 1.6 | < 0.1 to 5.8 |
| Davis et al. (2001a) | mangrove | Florida | 0.1 to 5.2 | 0.1 to 5.8 |
| Davis et al. (2001b) | mangrove | Florida | 0.1 to 6.3 | 0.2 to 5.8 |
| Ohowa et al. (1997) | mangrove | Kenya | 0.3 to 3 | 0.2 to 8 |
| Rivera-Monroy et al. (1995) | mangrove | Mexico | 1.1 to 51.7 | 0.2 to 4.9 |
| Rivera et al. (2007) | mangrove | Florida | < 0.1 to 1.4 | 1.8 to 11.8 |
| Eyre and Balls (1999) | estuary main channel | Australia | < 0.1 to 4 | < 0.1 to 7 |
| Eyre (1994) | estuary main channel | Australia | — | 2.3 to 31.2 |

**Figure 11:**
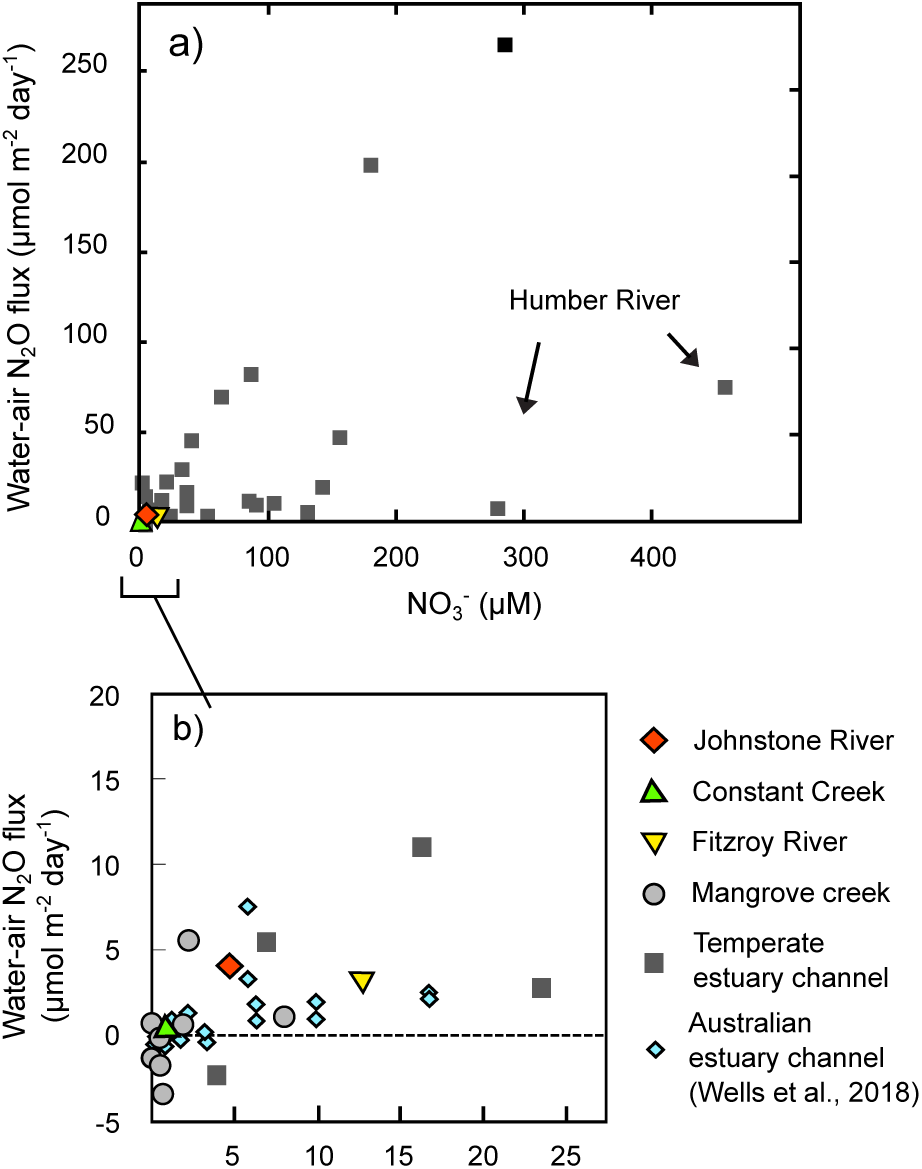
(a) Average annual NO_3_ ^−^ concentration (*µ*mol L^−1^) vs N_2_O water-air flux (*µ*mol m^−2^ day^−1^; modified from (Murray et al., 2015). (b) Data from the Johnstone River Estuary, Constant Creek, and the Fitzroy River Estuary, plotted against data presented in (Murray et al., 2015) along with results from several mangrove creeks (Barnes et al., 2006; Maher et al., 2016; Murray et al., 2018) and southeast Queensland estuaries (Wells et al., 2018).

The ratio of N_2_O emissions to total DIN load was quite low (0.02 – 0.09%), except for the Fitzroy dry season (1.12%). For comparison, N_2_O emissions accounted for 0.26% of DIN load in the Changjiang River in China (Zhang et al., 2010), about 0.29% of TN in the Colne River (Robinson et al., 1998), and 0.3% of DIN is assumed to be converted to N_2_O in estuaries in some global models of coastal N dynamics (Kroeze and Seitzinger, 1998). However, the ratio between DIN load and N_2_O emissions, referred to as the emissions factor, can vary significantly and lower values are common. In a review of emissions factors in rivers, Hu et al. (2016) found a range of values from 0.003% to 7.13%, with a median of 0.15%. They found that the emissions factor increases as freshwater discharge declines, which might explain why the value was so high in the Fitzroy River during the dry season (1.12%), when discharge was low (Figure 2c) and residence time was high (Figure 5).

## 6. Conclusions

In the Johnstone River, N_2_O came from a freshwater source upstream of the sampled estuary, with the highest contribution of N_2_O coming from the South Johnstone River. The N_2_O concentration of freshwater was similar in both the wet and the dry season, but the amount of freshwater delivered to the estuary in the wet season was much greater. In both seasons, the freshwater residence time was low enough that DIN and dissolved N_2_O moved through the estuary with little evidence of in-situ production or consumption. Apart from the N_2_O emitted through water-air exchange, dissolved N_2_O was largely exported to the ocean.

The Fitzroy River was affected by wastewater input from the sewage treatment plants near Rockhampton. This was most obvious in the dry season, when freshwater input was low, and led to a stronger production and water-air flux of N_2_O. The residence time was long, compared with the Johnstone River; Excess N_2_O from the riverine source was completely lost to the atmosphere and additional N_2_O from in-situ processes and downstream tributaries was added along the length of the transect.

Constant Creek was the site of N_2_O consumption in the wet season, due to the unusually low concentrations of DIN in the water-column. In the dry season there was a slight N_2_O production in some offshore shoals, but the N_2_O concentration and fluxes in the creek itself remained very low. Comparing these results with previous studies, it seems that a NO_3_ ^−^ concentration of less than 5 *µ*mol L^−1^ is conducive to a negative water-air N_2_O flux.

## Appendix

**Table A1:** The parameterization equations used to calculate gas transfer k-values (in cm s^−1^) from U_10_ (wind speed adjusted to 10 m altitude, in m s^−1^)

| Citation | Equation |
| --- | --- |
| Clark et al. (1995) | $k = (2 + 0.24 * U_{10}^2) * (Sc/600)^{-0.5}$ |
| Wanninkhof (2014) | $k = 0.251 * U_{10}^2 * (Sc/600)^{-0.5}$ |
| Liss and Merlivat (1986) | $\begin{cases} 0.17 * U_{10} * (Sc/600)^{-2/3} & \text{If } U_{10} \leq 3.6 \\ (2.85 * U_{10} - 9.65) * (Sc/600)^{-0.5} & \text{If } 3.6 > U_{10} > 13 \\ (0.59 * U_{10} - 49.3) * (Sc/600)^{-0.5} & \text{If } U_{10} > 13 \end{cases}$ |
| Ro and Hunt (2006) | $k = 5.9 * U_{10}^{1.81} * (Sc/600)^{-0.5}$ |
| Nightingale et al. (2000) | $k = (0.222 * U_{10}^2 + 0.333) * U_{10} * (Sc/600)^{-0.5}$ |
| Raymond and Cole (2001) | $k = 1.91 * e^{0.35 * U_{10}} * (Sc/600)^{-0.5}$ |

**Table A2:** Calculated k-values using 6 different parameterisations. The mean of all k-values is in parentheses.

| Location/season | k value ($\text{cm s}^{-1}$) | | | | | |
| --- | --- | --- | --- | --- | --- | --- |
|  | C 95 | W 14 | L&M 86 | R&H 06 | N 2000 | R&C 01 |
| Johnstone River wet season | 2.2–12.6<br>(5.8) | 0–11.3<br>(4) | 0–1.2<br>(0.7) | 0–7.3<br>(2.8) | 0–11.9<br>(4.6) | 2.1–18.8<br>(7.7) |
| Johnstone River dry season | 2.2–12.2<br>(5.4) | 0–10.8<br>(3.4) | 0–1.2<br>(0.6) | 0–7.1<br>(2.4) | 0–11.4<br>(4) | 2.1–17.8<br>(7.1) |
| Constant Creek wet season | 4.7–12.6<br>(8.5) | 2.8–11.4<br>(6.9) | – 0.8–9<br>(4.6) | 2.1–7.4<br>(4.6) | 3.5–11.9<br>(7.6) | 6.2–18.8<br>(11.6) |
| Constant Creek dry season | 8–18.5<br>(13.6) | 6.5–17.9<br>(12.5) | 4.3–14<br>(9.9) | 4.4–11.1<br>(8) | 7.2–18<br>(12.9) | 10.9–33.2<br>(21.3) |
| Fitzroy River wet season | 4.8–34.6<br>(16.9) | 2.8–35.3<br>(16) | – 0.9–24.7<br>(12.2) | 2.1–20.7<br>(10) | 3.6–34<br>(16.2) | 6.3–91.4<br>(29.8) |
| Fitzroy River dry season | 2.5–18.6<br>(7.7) | 0.5–18.3<br>(6.3) | – 5.6–14<br>(3.9) | 0.4–11.2<br>(4.2) | 0.9–18.2<br>(6.9) | 3.1–36<br>(11) |

**Figure A1:**
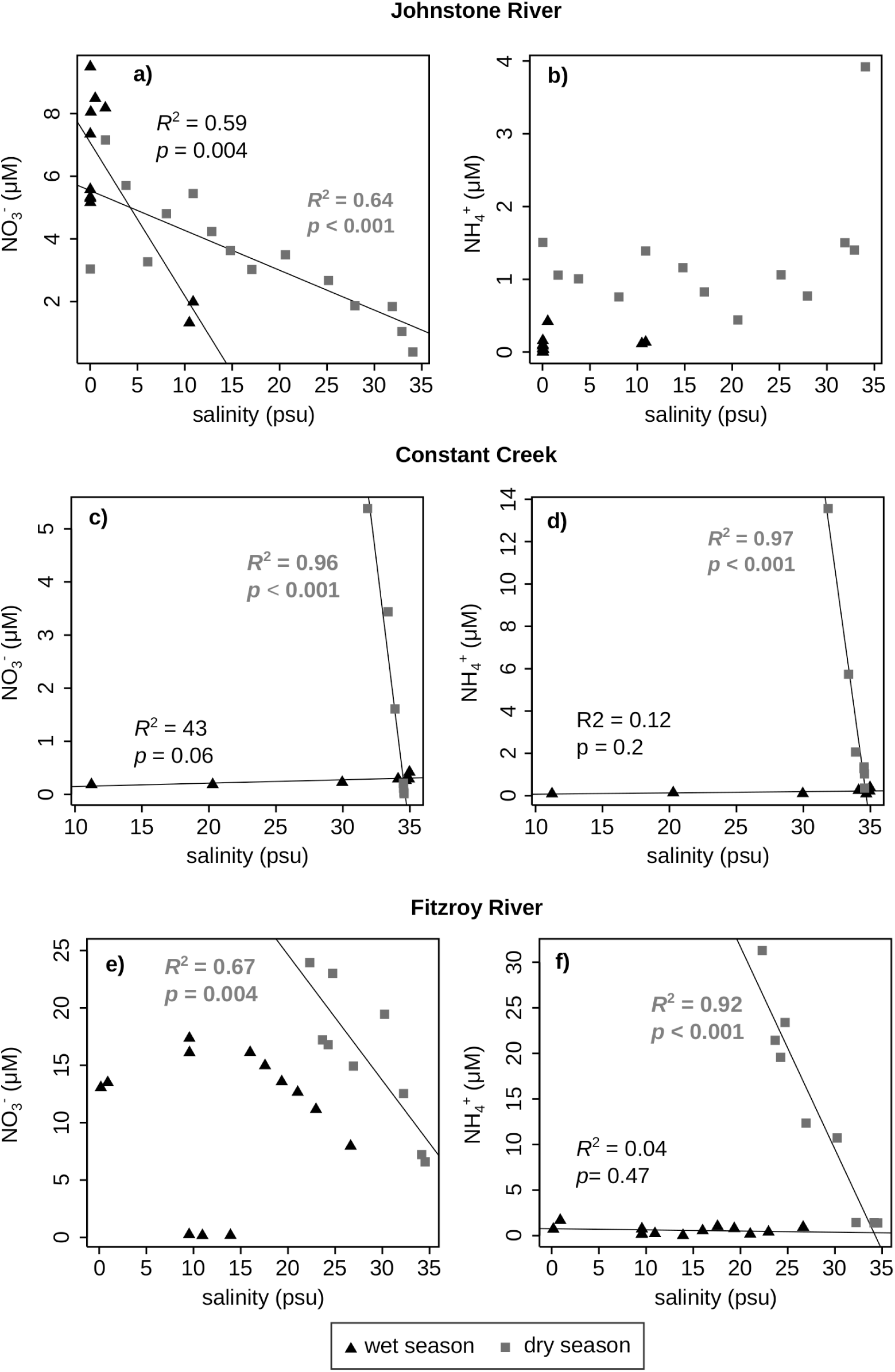
Salinity vs NO_3_ ^−^ and NH_4_^+^ during the wet and dry season in the Johnstone River Estuary, Constant Creek, and the Fitzroy River Estuary.

**Figure A2:**
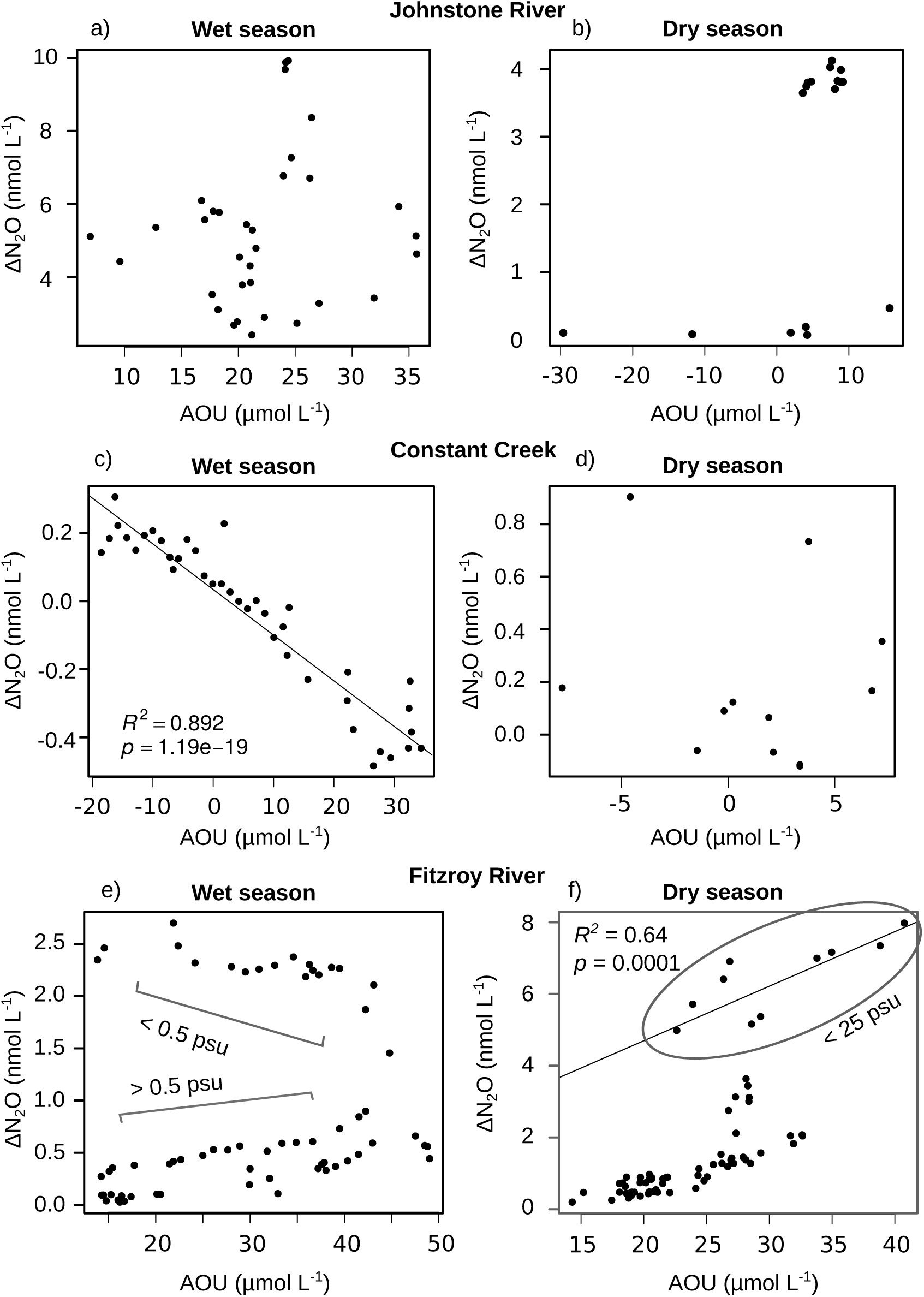
AOU vs ΔN_2_O during the wet and dry season in the Johnstone River Estuary, Constant Creek, and the Fitzroy River Estuary.

**Table A3:**
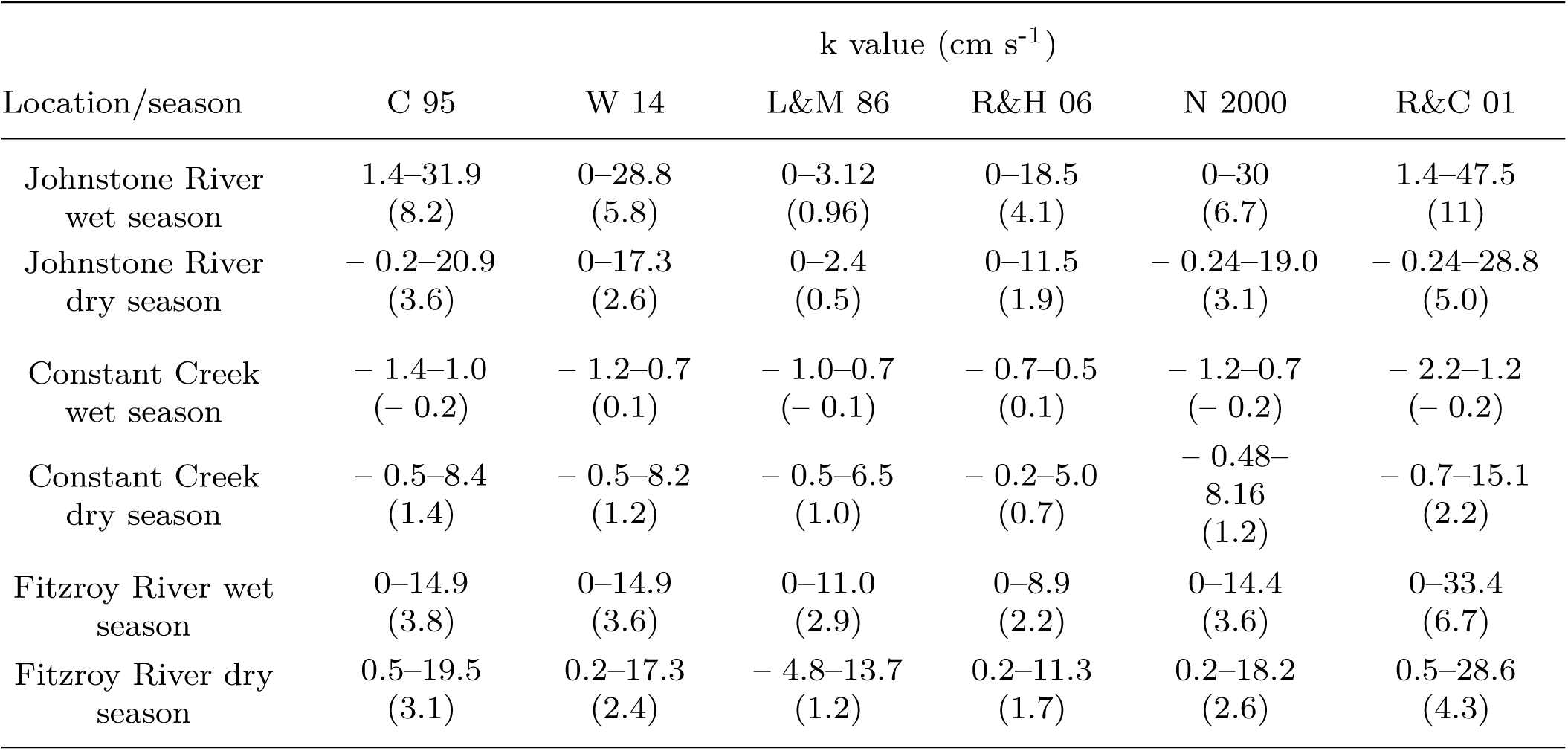
Results of calculations for water-air N_2_O flux, using 6 different parameterisations. The mean flux is in parentheses.

| Location/season | k value (cm s<sup>-1</sup>) |  |  |  |  |  |
| --- | --- | --- | --- | --- | --- | --- |
|  | C 95 | W 14 | L&M 86 | R&H 06 | N 2000 | R&C 01 |
| Johnstone River<br>wet season | 1.4–31.9<br>(8.2) | 0–28.8<br>(5.8) | 0–3.12<br>(0.96) | 0–18.5<br>(4.1) | 0–30<br>(6.7) | 1.4–47.5<br>(11) |
| Johnstone River<br>dry season | – 0.2–20.9<br>(3.6) | 0–17.3<br>(2.6) | 0–2.4<br>(0.5) | 0–11.5<br>(1.9) | – 0.24–19.0<br>(3.1) | – 0.24–28.8<br>(5.0) |
| Constant Creek<br>wet season | – 1.4–1.0<br>(– 0.2) | – 1.2–0.7<br>(0.1) | – 1.0–0.7<br>(– 0.1) | – 0.7–0.5<br>(0.1) | – 1.2–0.7<br>(– 0.2) | – 2.2–1.2<br>(– 0.2) |
| Constant Creek<br>dry season | – 0.5–8.4<br>(1.4) | – 0.5–8.2<br>(1.2) | – 0.5–6.5<br>(1.0) | – 0.2–5.0<br>(0.7) | – 0.48–<br>8.16<br>(1.2) | – 0.7–15.1<br>(2.2) |
| Fitzroy River wet<br>season | 0–14.9<br>(3.8) | 0–14.9<br>(3.6) | 0–11.0<br>(2.9) | 0–8.9<br>(2.2) | 0–14.4<br>(3.6) | 0–33.4<br>(6.7) |
| Fitzroy River dry<br>season | 0.5–19.5<br>(3.1) | 0.2–17.3<br>(2.4) | – 4.8–13.7<br>(1.2) | 0.2–11.3<br>(1.7) | 0.2–18.2<br>(2.6) | 0.5–28.6<br>(4.3) |

